# Cassette recombination dynamics within chromosomal integrons are regulated by toxin–antitoxin systems

**DOI:** 10.1101/2022.08.03.502626

**Authors:** Egill Richard, Baptiste Darracq, Eloi Littner, Claire Vit, Clémence Whiteway, Julia Bos, Florian Fournes, Geneviève Garriss, Valentin Conte, Delphine Lapaillerie, Vincent Parissi, François Rousset, Ole Skovgaard, David Bikard, Eduardo P. C. Rocha, Didier Mazel, Céline Loot

## Abstract

Integrons are adaptive bacterial devices that rearrange promoter less gene cassettes into variable ordered arrays under stress conditions, to sample combinatorial phenotypic diversity. Chromosomal integrons often carry hundreds of silent gene cassettes, with integrase-mediated recombination leading to rampant DNA excision and integration, posing a potential threat to genome integrity. How this activity is regulated and controlled, particularly through selective pressures, to maintain such large cassette arrays is unknown. Here we show a key role of promoter-containing toxin–antitoxin (TA) cassettes as abortive systems that kill the cell when the overall cassette excision rate is too high. These results highlight the importance of TA cassettes regulating the cassette recombination dynamics and provide insight into the evolution and success of integrons in bacterial genomes.

**Teaser:** The accumulation of cassette functions in integrons is ensured by toxin–antitoxin systems which kill the cell when the cassette excision rate is too high.

## Introduction

Integrons are bacterial recombination systems that are capable of capturing, stockpiling, and reordering mobile elements called cassettes, to control their expression. They were originally found to be the genetic systems responsible for the acquisition of antimicrobial resistance determinants in some mobile elements (*1*). Integrons contain a stable platform and a variable cassette array (Fig 1a). The stable platform is defined by i) the integrase gene (*intI*) under the control of its promoter P_int_, ii) the *attI* integration site and iii) the cassette P_C_ promoter. The cassettes in the array are generally composed of a promoter less gene associated to a recombination site called *attC*. Only the first few cassettes (those closest to the P_C_ promoter) can be expressed, while the rest represent silent and valuable functions for the cell (*2*). These cassettes can be excised and then re-integrated at the *attI* integration site and therefore become expressed (*3*). This cassette shuffling ensures a combinatorial phenotypic diversity which allows bacteria to screen for the set of functions that would optimize its survival in a given environment. Remarkably, we previously found that the integrase, which catalyzes the cassette shuffling, is only expressed in stress conditions (*4*). For this reason, integrons are described as “on demand” adaptation systems (*5, 6*).

**Figure 1:**
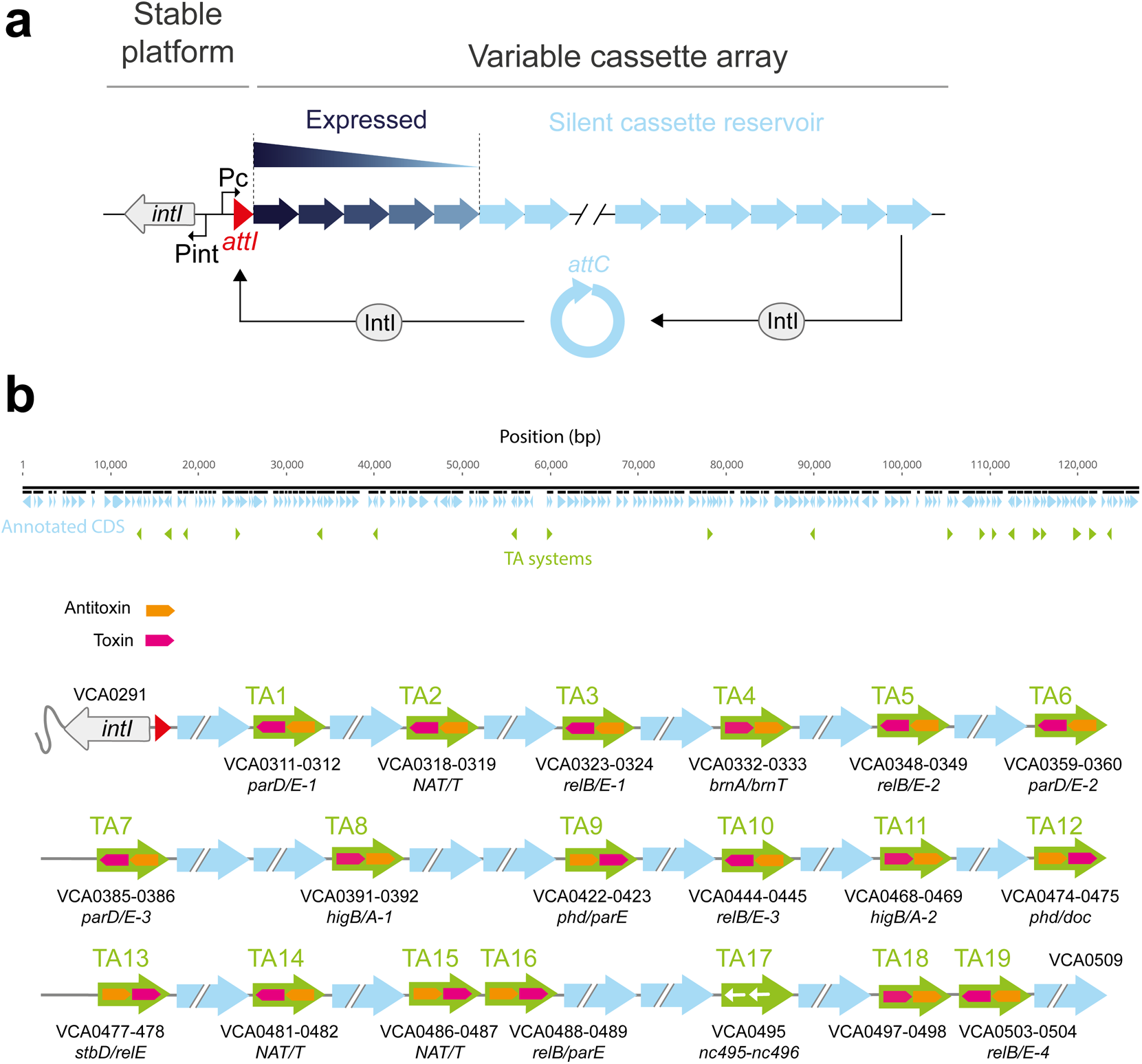
The integron system and the toxin–antitoxin cassettes. **a.** Schematic representation of the integron. The stable platform consists of a gene coding for the integrase (*intI*, in yellow) and its promoter Pint, the cassette insertion site *attI* (in red) and the cassette promoter P_C_ driving the expression of the downstream cassettes along an expression gradient. Cassettes are displayed as arrows where the tip represent the *attC* site. Their color intensity represents their level of expression from dark blue (highly expressed) to light blue (not expressed). Non-expressed cassettes can be excised through an intramolecular *attC* × *attC* reaction and be reintegrated in 1^st^ position near the P_C_ promoter through an intermolecular *attC* × *attI* reaction and become expressed. The combination of excision and integration is referred to as cassette shuffling. **b.** Representation of the SCI of *V. cholerae* and the repartition of the various toxin–antitoxin (TA) systems numbered from 1 to 19. A detail of the TA systems (name, orientation in the array, order of toxin/antitoxin) is represented below. NAT/T means N-acetyltransferase/transcription factor.

The integron integrase (IntI) has a very singular place among the broad family of the tyrosine recombinases (*7*). While it recognizes the *attI* site under its double stranded (ds) form through its primary sequence, the *attC* sites are recombined as a single stranded (ss) form (*8, 9*). Although both the bottom and the top strands of an *attC* site can form a secondary structure, the recombination of the bottom strand (*bs*) is about 10^3^ more efficient than that of the top (*9*). This ensures a correct orientation of the cassette upon integration at the *attI* site, allowing its expression by the P_C_ promoter (*10*).

We distinguish two types of integrons: Mobile Integrons (MIs) carried by conjugative plasmids and mobilizable among bacteria, and Sedentary Chromosomal Integrons (SCIs). In addition to their sedentary nature, the main feature that distinguishes SCIs from MIs is the size of their cassette array. While MIs do not typically store more than 10 cassettes, SCIs often contain dozens of cassettes — with the largest SCI from *Vibrio vulnificus* encoding 301 — thus encompassing a substantial fraction of their host genomes (*11*). SCI cassettes constitute an almost infinite variety of functions extending far beyond antibiotic resistance and linked to bacterial key adaptive functions (*12*). The high genetic capacitance of SCIs allows them to serve as a repository of cassettes for MIs to compile the most relevant repertoire of cassettes in a specific environment (*6, 13*). The remarkable adaptive success of the integron system is partly due to this intricate connection between SCIs and MIs. Despite all that, twenty-five years after their discovery, SCIs remain enigmatic genetic structures. We still ignore how such structures can accumulate so many silent cassettes and be maintained in bacteria to provide such an adaptive device for future stresses. Among the cassettes of SCIs, a distinct type is notable, encoding toxin–antitoxin (TA) systems and containing their own promoter (*14-17*). TA systems encode a stable toxin that inhibits an essential cellular process and an unstable antitoxin that counteracts its cognate toxin. When the expression of the gene pair is stopped, the antitoxin is degraded, resulting in cell death or growth arrest (*18*). The presence of toxin–antitoxin systems within SCI arrays likely prevents cassette deletion through large chromosomal rearrangements thus maintaining the array (*14, 15*). Nevertheless, this sole factor is not sufficient to understand how SCIs are kept so massive. Indeed, in stress conditions, when the integrase is expressed, the cassette excision rate must be tightly regulated. To ensure integron cassette accumulation, the cassette excision rate must be kept below the integration rate, and to maintain the large integrons, the two rates must be at least similar. An imbalance in favor of the excision rate would lead *in fine* to the loss of most silent cassettes. Here, directly linked to their cassette nature, we demonstrated that the TAs play an additional role regulating the cassette recombination dynamics. We used as model the *Vibrio cholerae* N16961 strain responsible for the ongoing 7^th^ cholera pandemic and which contain a particularly massive SCI (∼130 kb and 179 cassettes (*19*)). By developing a genetic assay to measure the SCI cassette dynamics at the population level, we demonstrate that, the cassette excision rate is dramatically increased all along the SCI when we “artificially” inverted it on-site. We show that the inversion of the SCI is also associated to a strong growth defect correlated with a high cell mortality rate in presence of a functional integrase only. By successfully inactivating all the 19 TA cassettes within the inverted SCI (*16, 17*) (Fig 1b), using highly multiplexed CRISPR base editing, we demonstrate that these cassettes are in a large part responsible for the observed mortality phenotype.

Our results highlight a new role for TA cassettes via “abortive cassette excision” depending on the integrase. Under conditions where the cassette excision rate is high and could compromise the maintenance of large SCIs, TA cassettes are also excised at a high rate as individual cassettes, thus killing cells. Through this, TAs exert a selective pressure that drives integrons to adopt a configuration in which the excision rate of their cassettes is kept low. This has probably led to the preferential orientation of SCI with respect to the replication fork that we previously observed (*13*), and which limits the rate of excision. The discovery of this new type of regulation in integrons adds to the already known regulatory network. It enables silent cassettes to be accumulated and maintained, making them available to produce combinatorial phenotypic diversity and respond to future stresses.

## Results

### SCIs are specifically enriched in toxin–antitoxin systems

To better understand the role of the TAs as cassettes in SCI arrays and to assess whether SCI size and TA number are correlated, we systematically explored all TA carried by SCIs. We used IntegronFinder 2.0 (*20*) to detect all of the sedentary integrons in complete genomes of the RefSeq NCBI database. We identified 398 genomes with at least one SCI (defined as an integron comprising strictly more than 10 cassettes, as suggested in (*20*)). In each of these genomes, we screened TA systems with TASmania (*21*). Then, we compared the number of toxins and antitoxins detected in the SCIs with the total number of cassettes in each of the 398 genomes. We found a positive correlation between the two, indicating that larger SCIs have more TA cassettes (Fig 2a). To assess the specific enrichment of TAs in SCIs when compared to the rest of the genome, we performed in each isolate a contingency table analysis. Out of the 398 isolates, 246 verified the hypothesis of a TA enrichment within SCI (p<0.05, Fisher test with Benjamini-Hochberg correction). Of the remaining isolates, 87 genomes lack genes encoding toxins or antitoxins and 65 encode at least one of them but do not show significant enrichment. As most of the genomes with a significant over-representation of TAs in SCIs belonged to the *Vibrio* genus (Fig 2a), we decided to analyze them thoroughly. Among the 280 *Vibrio* genomes, 225 verified the hypothesis of a TA enrichment within SCI (Fig 2b). This may reflect a particularly strong effect in this genus, but we cannot exclude that it simply results from other SCIs having too few cassettes to provide a statistically significant signal of over-representation. We studied the patterns of this over-representation in function of the phylogeny of *Vibrio*. We observe that the enrichment signal is scattered across the tree, highlighting the important role of TAs in a broad range of *Vibrio* species (Fig 2c). This association holds true with the very stringent Bonferroni correction, although with a less sparse distribution on the phylogenetic tree, centered around a clade comprising *V. cholerae* (Fig S1). Interestingly, in this species, some TA systems (in the parDE family) are conserved in most isolates (82/86). This conservation may suggest a long period of co-evolution of integrons and TA systems, at least in *V. cholerae*. Although it does not exclude the regular turnover of some other TA systems, TA cassettes may have contributed to the evolution of SCIs to a large extent.

**Figure 2:**
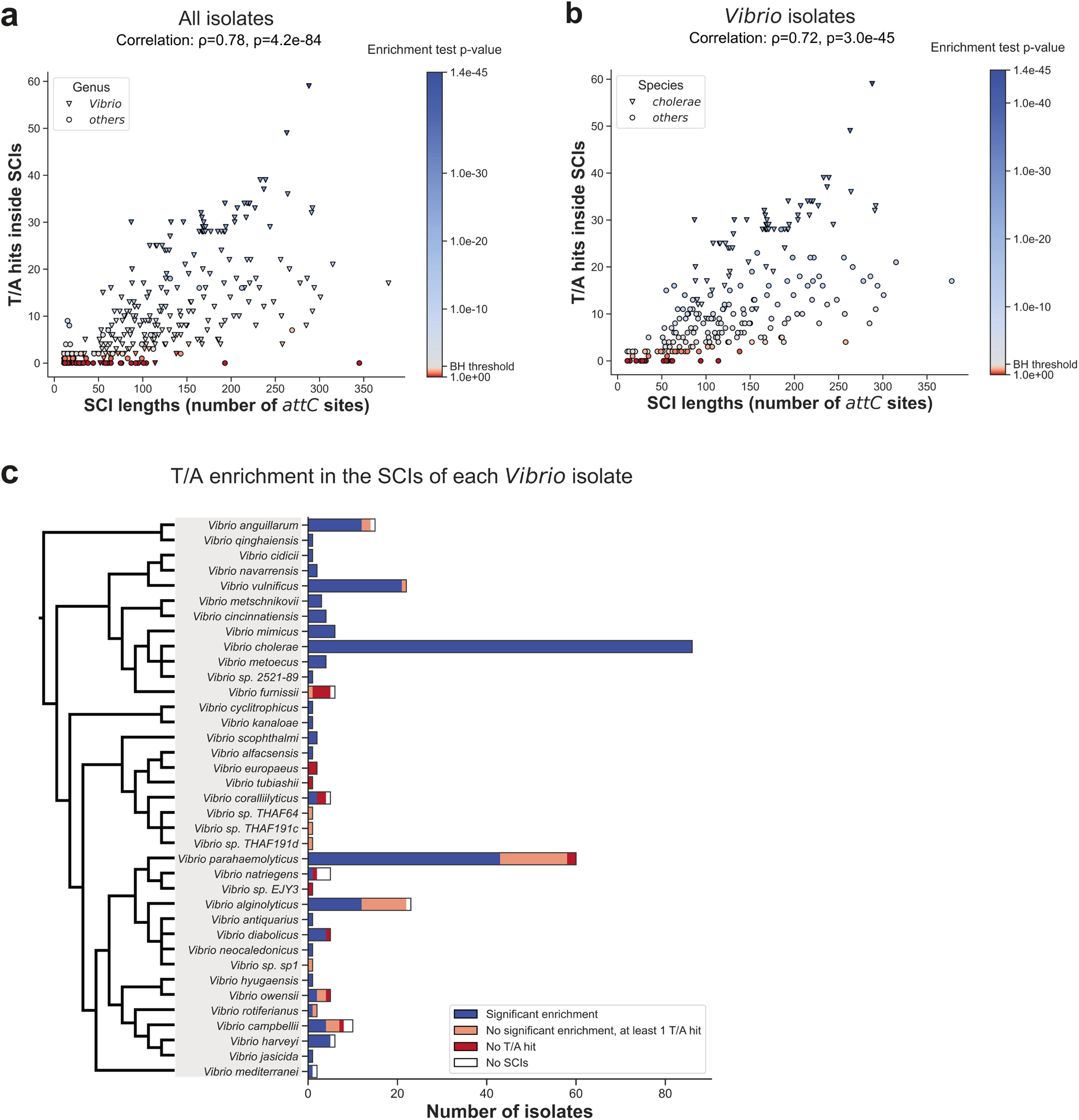
Toxin–antitoxin landscape in SCIs. **a, b.** Each point represents a single NCBI RefSeq isolate harboring at least one SCI (>10 *attC* sites). The x-axis displays the total number of *attC* sites in the (potentially multiple) SCI of the isolate. The y-axis exhibits the total number of TASmania toxin and antitoxin HMM hits in the SCI of the isolate (e-value<10^-3^). The Spearman nonparametric rank correlation coefficient is shown at the top of the graph, together with its significance p-value. Point hue indicates the p-value of Fisher’s test for TASmania hits enrichment in SCI compared to the rest of the genome (scale on the right of the graph, significant enrichment in blue). The Benjamini-Hochberg correction for multiple testing was applied (see below for the corresponding computed thresholds) **a.** All the isolates harboring at least one SCI are displayed. The shape of each point on the graph represents whether the isolate belongs to the *Vibrio* genus (triangles) according to the NCBI taxonomy or to another genus (circles). Benjamini-Hochberg correction threshold: 0.031. **b.** Same graph as a, but only with isolates belonging to the *Vibrio* genus. The shape of each point on the graph represents whether the isolate belongs to the *Vibrio cholerae* species (triangle) or to another species (round). Benjamini-Hochberg correction threshold: 0.036. **c.** Distribution of *Vibrio* isolates by species according to the toxin/antitoxin enrichment inside the SCI they harbor. The cladogram used to group species was adapted from the phylogeny reconstructed by Sawabe and coll (*22*). T: toxin, A: antitoxin BH: Benjamini-Hochberg correction

### The inversion of the SCI in *V. cholerae* dramatically increases the cassette excision rate in the array

Evolutionary factors have enabled the integron cassette accumulation and maintenance to lead to the emergence of massive and stable chromosomal integrons. To specify these factors and determine if TA cassettes have a role in this evolution process, we decided to render the cassette array of large integrons unstable and determine how these “genetically modified” integron structures can be counter-selected. To do this, we inverted the *V. cholerae* SCI on place (SCI Inv strain) and re-inverted it back to its original orientation (SCI Reinv strain) (Fig 3a, Fig S2 and Materials and Methods). Indeed, we previously establish that when the recombinogenic *bs* of *attC* sites are located on the lagging strand template, where discontinuous replication leaves single-strand gaps, this favors *attC* site folding and cassette excision (*13, 23*). We therefore hypothesized that the natural SCI orientation, where the *bs* of *attC* sites are located on the leading strand template, disfavors cassette excision events and that the SCI inversion would be a way to make the cassette array unstable.

**Figure 3:**
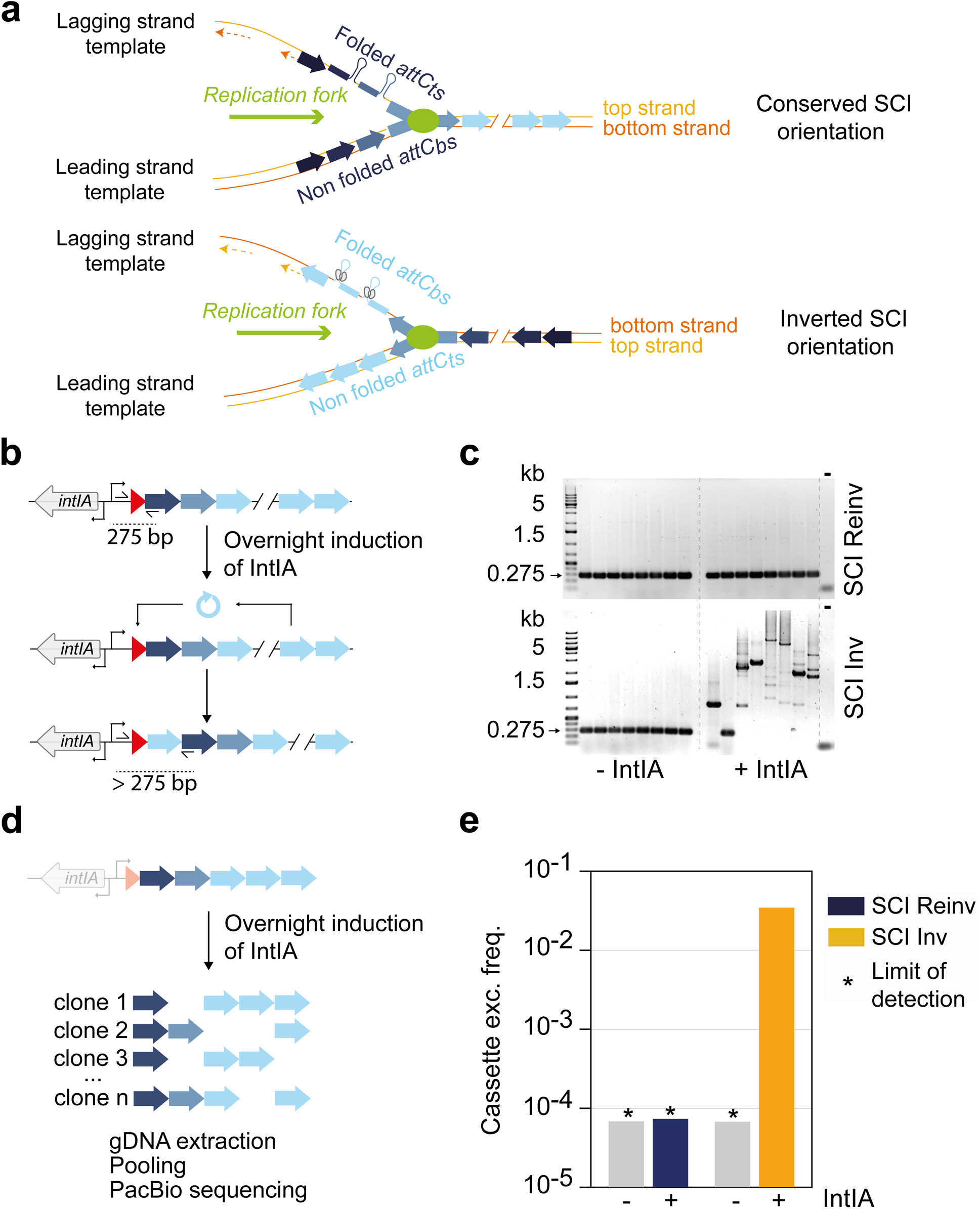
Cassette shuffling and excision assays in the SCI inverted and reinverted *V. cholerae* strains. **a.** Mechanistic insight on the issue of integron orientation. An array of cassettes is represented while it is replicated. In the lagging strand, Okazaki fragments are represented as dotted lines, leaving stretches of ssDNA on the correspondent template. In their conserved orientation, SCIs recombinogenic *attCbs* are carried by the continuously replicated leading strand template which supposedly limits their structuration. The structured *attCts* are not recombined by the integrase. In the inverted orientation their more frequent structuration is expected to lead to increased binding of the integrase and higher recombination frequencies. **b.** Schematic representation of the assay designed to detect cassette shuffling. **c.** PCR analysis of the cassettes shuffled in 1^st^ position in individual clones in presence (+IntIA) or in absence of the integrase (-IntIA) for one overnight. The expected PCR product in absence of shuffling is 275 bp. Longer bands are evidence shuffling events. Multiple bands may come from the random slippage of the DNA polymerase in presence of highly repeated sequences (multiple *attC* sites in one amplicon). **d.** Schematic representation of the assay designed to detect cassette excision. A strain of *V. cholerae ΔattIA* is used to initiate a culture composed of clones experiencing various cassette excision events. Clones from this heterogeneous population are represented (from 1 to n) each with different cassettes missing. **e.** Cassette excision frequency in the array of the SCI. The cassette excision frequency is calculated for the SCI inverted strain expressing the integrase (see Materials and Methods). No excision event could be detected in the other conditions; hence the limit of detection (*) is represented instead. The grey bars represent the frequencies obtained in absence of integrase in the SCI Inv and SCI Reinv strains.

We first monitored the effect of SCI inversion on cassette shuffling by cultivating overnight *V. cholerae* with the SCI in both orientations, in presence or absence of IntIA (Fig 3b). A PCR was then performed on independent clones with primers located in the *attIA* integration site and in the first cassette of the array (VCA0292) with an expected size of 275 bp in absence of shuffling (Fig 3b). Very few shuffling events could be detected in the SCI Reinv clones (3/48), whereas we observed large amplicons in 46/48 of the SCI Inv clones, testifying to the insertion of multiple cassettes at the *attIA* site (Fig 3c). Performing our classical conjugation assay to deliver cassettes in *V. cholerae* recipient strains (*3*), we obtained a constant integration rate of these cassettes at the *attIA* site regardless of the SCI orientation (Fig S3). Our results therefore demonstrate that SCI inversion results in a substantial increase in cassette excision while the integration rate remains stable.

To monitor the cassette excision rate at the population level in strains with SCI in both orientations, we designed an SCI-wide cassette excision assay. Like the previous shuffling assay, we grew *V. cholerae* strains overnight in the absence or presence of integrase, but this time in the absence of the *attIA* site to focus solely on excision events. To assess cassette rearrangements along the array, we used long-read sequencing (Materials and Methods, Fig 3d). Each read captured the state of a segment of the SCI in a single cell and allowed an easy detection of cassette deletion on this segment. For each population, a total of ∼80000 reads with an average length of ∼6.4 kb was produced. For the *V. cholerae* population corresponding to the SCI inverted strain expressing integrase, we identified 478 cassette excision events, encompassing 120 cassettes where excision was observed at least once leading to a cassette excision rate of 3 × 10^-2^ (Fig 3e). For the populations of strains with the SCI in the native orientation with and without integrase and, in the strain with an inverted SCI in absence of the integrase, the excision rates are below 7 × 10^-5^ (*i.e.*, the limit of detection, Fig 3e and Materials and methods). These results reveal a direct link between the inversion of the SCI and a highly increased cassette excision frequency.

### The inversion of the SCI in *V. cholerae* leads to a growth defect in presence of a functional integrase

Genome rearrangements are not very rare (*24, 25*), and if a single inversion is sufficient to increase the excision cassette rate and empty a SCI of all its cassettes in a few generations, then the massive and silent structures that are SCIs would probably not exist. We reasoned that there may be counter-selective forces in situations in which the cassette array is unstable, preventing, for example, a spontaneous inversion of the SCI. We set out to measure growth of the SCI Inv strain (Fig 4a). While we did not observe any growth defect in the SCI Inv strain in absence of integrase, we found an important growth defect upon expression of the integrase in this strain compared to the SCI Reinv strain (Fig 4a). Growth rate is ∼35% lower in the SCI Inv strain expressing the integrase compared to the controls (Fig 4b). We observed the same growth defect while inducing the endogenous integrase through the SOS response using sub-lethal concentrations of ciprofloxacin (Fig 4c), confirming our previous conclusions in this more natural setting. To further evaluate the cost of the inversion of the SCI, we set out a competition experiment where SCI Inv and Reinv strains were co-cultivated for 24h starting with an initial 1:1 ratio (Fig 4d). While no substantial deviation to the original ratio between SCI Inv and Reinv strains could be observed in absence of integrase, the expression of the integrase in those strains quickly led to a strong disadvantage of the SCI Inv strain compared to the SCI Reinv. This confirms the high fitness cost of the inversion of the SCI of *V. cholerae* in presence of the integrase. Expressing a catalytically inactive integrase that binds to the *attC_bs_* (Fig S4) without cleaving it did not produce any growth defect (Fig 4e), ruling out a scenario where the mere binding of more accessible *attC_bs_* in the SCI Inv strain could explain the growth defect. Hence, the cleavage activity of the integrase was necessary to induce a growth defect in the SCI Inv. Yet, the integrase can cleave both *attIA* and *attC* sites so to determine if it was the increased shuffling or the mere cassette excision that imposed such a strong cost of SCI inversion, we performed growth curve in cells lacking the *attIA* site (Fig 4f). We still observed a growth defect, hence confirming that it is the increased cassette excision rate that explained the growth defect of the SCI Inv strain.

**Figure 4:**
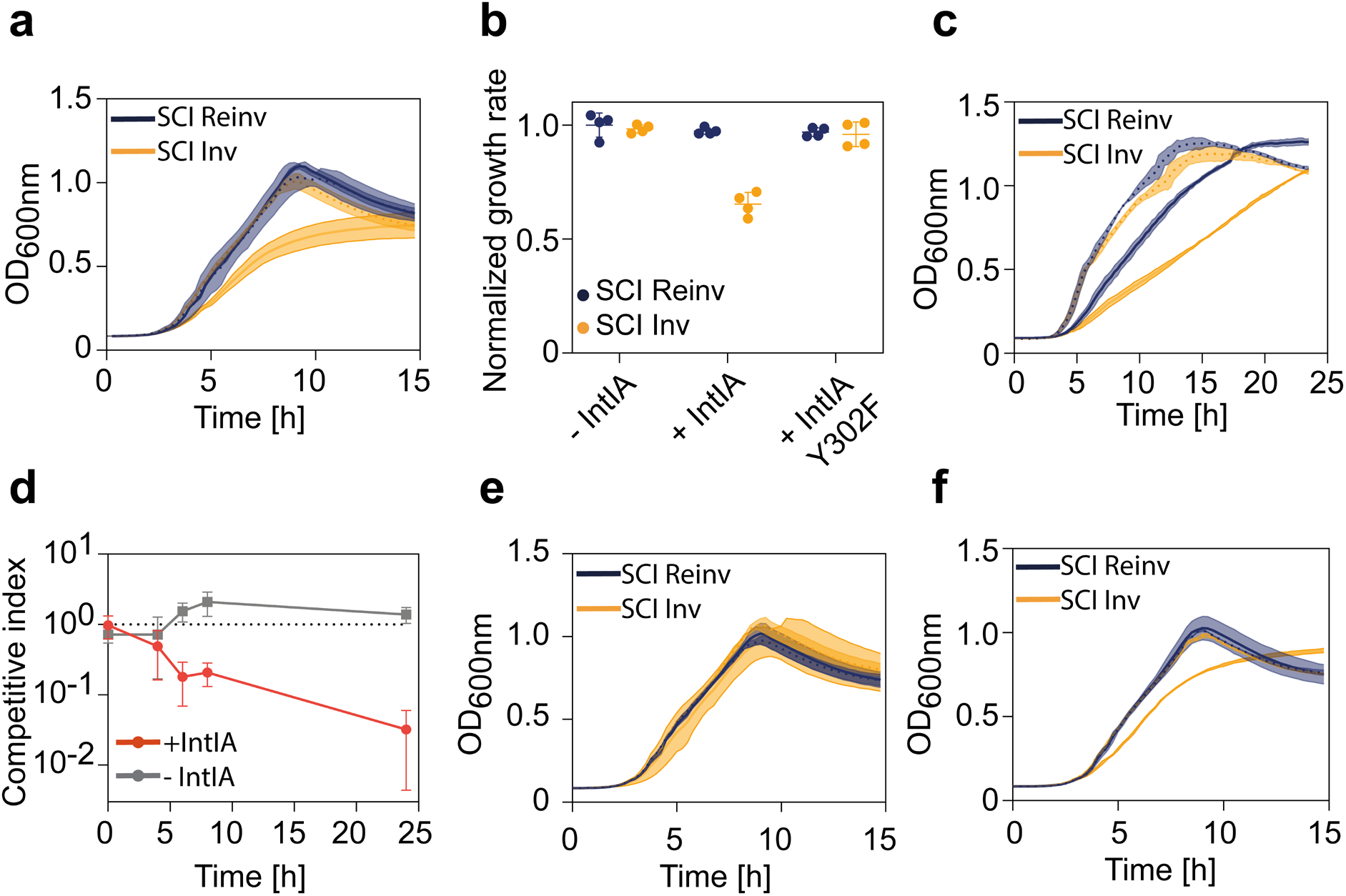
Growth of the SCI inverted and reinverted *V. cholerae* strains. **a.** Growth curve of SCI Inv (orange) or SCI Reinv (blue) *V. cholerae* strains in presence (full lines) or in absence of IntIA (dotted lines). Curve corresponds to the mean of four biological replicates, each with an average of at least two technical replicates, and the shade corresponds to the standard errors at each timepoint (same for all growth curves). **b.** Growth rates of the SCI Inv and SCI Reinv strains normalized by the mean growth rate of the SCI Reinv strain in absence of IntIA. The normalized growth rate of each strain respectively in absence of integrase (- IntIA) and, in presence of the WT (+ IntIA) or the catalytically inactive integrase (+ IntIA Y302F) are represented. **c.** Growth curve of the SCI Inv and Reinv strains in presence (full lines) or in absence (dotted lines) of a sub-inhibitory concentration of ciprofloxacin (10 ng/ml) inducing the endogenous IntIA expression. **d.** Competitive index of the SCI Inv strain compared to the Reinv strain in absence (gray) or in presence of IntIA (red) in function of time (24h co-cultures). An index of 1 represents a ratio of 1:1 of the two strains in the mix. The lower the index, the lower the ratio of SCI Inv compared to SCI Reinv. Index was calculated with three biological replicates for each time point and the means and standard errors are represented. **e.** Growth curve of the SCI Inv and Reinv strains in presence of the catalytically inactive IntIA (full lines) or in absence of IntIA (dotted lines). **f.** Growth curve of the SCI Inv and Reinv strains in presence (full lines) or in absence (dotted lines) of IntIA, this time in a strain lacking an *attIA* site.

### The growth defect in the SCI Inverted strain in presence of integrase is associated with increased cell death

The growth defect previously observed in the integrase-expressing SCI Inv strain could result from a slower division rate of each individual cell or a higher mortality rate at each generation. To discriminate between these two possibilities, we needed information at the single cell level. We therefore set out to observe in widefield microscopy live cells of our different strains growing on an agarose pad mimicking the liquid medium used for generating the growth curves. An exponentially growing culture of the SCI Inv and Reinv strains was diluted to display individual cells onto the agarose pads. We recorded the development of microcolonies for 170 min (Fig 5a, Movie S1a, S1b, S2a and S2b) and the number of cells per microcolony in the SCI Inv and Reinv strains expressing the integrase was determined as a measure of growth. We observed that the SCI Inv strain displayed increased heterogeneity in growth across microcolonies resulting in a broader distribution of microcolony sizes at the end of the run (from t=120 min, Fig 5b, Movie S2a and S2b). This reflects the fact that, whereas the SCI Reinv colonies generally follow an expected developmental trajectory, in the SCI Inv strain, many microcolonies stop growing after a few divisions or do not grow at all. Some events of cell lysis, characterized by sudden release by some cells of diffracting matter (*26*), could also be observed specifically in the SCI Inv strain (Fig 5c, Movie S2c). These observations favor a model where the expression of the integrase in the SCI Inv strain induces cell death in a part of the population.

**Figure 5:**
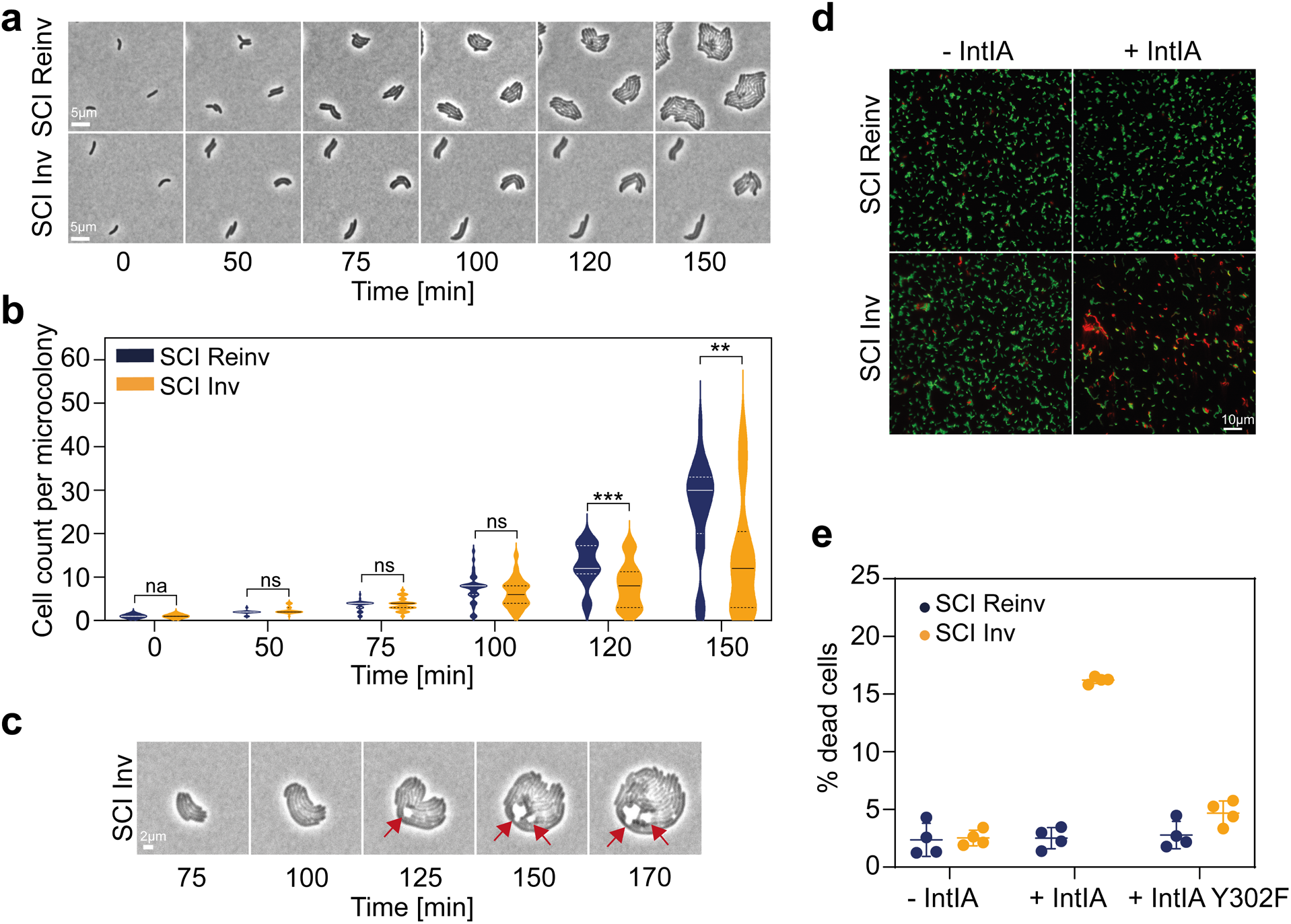
Cell viability of the SCI inverted and reinverted *V. cholerae* strains. **a.** Bright-field microscopy images taken from 170 min time-lapse series of live cells growing onto a MOPS agarose pad. A representative sample of 3 microcolonies of SCI Inv or SCI Reinv strains is shown. Scale bar is 5 µm. **b.** Microcolony growth of the SCI Inv and Reinv strains onto MOPS agarose pads. Violin plots show the distribution of the number of cells per microcolony (27 for SI Inv and 34 for SCI Reinv strains) tracked during 170 min. Median (full line) and quartiles (dotted line) are shown. Statistical comparisons (Mann-Whitney test) are as follow: (na. (not applicable), ns (not significant), *** *Pvalue < 0.001,* \**\* < 0.01*). **c.** Representative example of cell lysis events (red arrows) captured during the time-lapse experiments of live SCI Inv cells growing on MOPS agar pads and expressing the integrase. Lysis events were not observed in the SCI Reinv strain expressing the integrase. Scale bar is 2 µm. **d.** Fluorescence microscopy images resulting from the Live and Dead assay on the SCI Inv and Reinv strains expressing or not the integrase. Green fluorescence indicates live cells (SYTO-9) whereas red fluorescence (Propidium iodide (PI)) indicates dead cells. Scale bar is 10 µm. **e.** Quantification of cell death as a measure of cells positively stained with PI over the total number of cells counted (∼ 1000 cells per replicate). Four biological replicates, means and standard deviation are represented.

We tested this model in a live and dead assay where a differential coloration of the cells allows discrimination between living (green) and dead (red) cells in a population (Fig 5d). We observed a clear increase of the proportion of dead cells in culture of SCI Inv strain expressing the integrase compared to the control conditions. We quantified the cell mortality and observed a high percentage of dead cells (∼16%) in the SCI Inv strain expressing the integrase and a basal mortality rate (∼3-5%) in the other strains (Fig 5e). Interestingly, expression of a catalytically inactive integrase in the SCI Inv strain resulted to a mortality rate similar to the basal level (Fig 5e), consistent with the observation that was made in the previous growth measurement (Fig 4b and 4e). These data indicated that inversion of the SCI led to a high mortality rate that was dependent on the activity of the integrase.

### The fitness defect in the SCI Inverted strain is largely explained by the excision of toxin– antitoxin systems

The fact that SCI inversion results in increased cassette excision and that this is associated with increased growth arrest and cell death dependent on integrase catalytic activity led us to hypothesize that the increased excision of some cassettes could be deleterious for the cell. We hypothesized that the increased excision of the cassettes encoding TA systems, alongside all the non-expressed cassettes, could be at the origin of the increased growth arrest and cell death observed in the SCI Inv strain. Indeed, analysis of *attC* sequences indicate that TA systems are as likely to be excised as other cassettes (Materials and methods, Fig S5a). In addition, we observe excision of the TA-encoding cassettes when they are duplicated (Fig S5b), suggesting that the non-duplicated TA cassettes can also be excised but result in such a large fitness defect that we cannot observe their excision in our data set. By combining prior approaches (*27, 28*), we designed a CRISPR-based cytosine base editors which allowed us to inactivate almost all the TA cassettes of the array in one go. 12 guides were designed to target 15 toxins (3 of which are duplicated) such that a STOP codon was introduced, resulting in at least a one-third truncation of each toxin (Materials and methods, Fig S6). The remaining 4 toxins, for which we were unable to design a suitable guide, were inactivated by introducing a stop codon through the more classical allelic exchange method (*29*). It resulted in *V. cholerae* strains where all the TA within the SCI were inactivated (TAi).

We next assessed the effect of TA inactivation on growth of both SCI Inv and Reinv strains. In absence of integrase (Fig 6a), we do not observe substantial differences between all the strains whether the SCI is inverted or not. In presence of integrase, the growth defect characteristic of the SCI Inv strain appears to be largely rescued upon inactivation of the TA systems (Fig 6b). Performing a competition assay, we found again, in presence of integrase, the SCI Inv to be severely out competed by the SCI Reinv strain but this was alleviated in absence of functional TA systems (Fig 6c). Moreover, the SCI Inv TAi outcompetes the SCI Inv strain by more than a factor of three in presence of integrase (Fig 6d). Note that both TAi strains, compared to the *wt* ones, present a slightly impacted fitness (Fig 6d), probably due to off-target mutations consequently to the TA inactivation process. A cell viability assay and a quantification of dead cells showed that the improved fitness of the SCI Inv TAi strain in presence of integrase compared with the SCI Inv strain was associated with decreased cell death (Fig 6e, 6f, 6g and 6h). For more clarity, we reported in these figures, the previous results presented in the figure 5 (i.e. TA WT).

**Figure 6:**
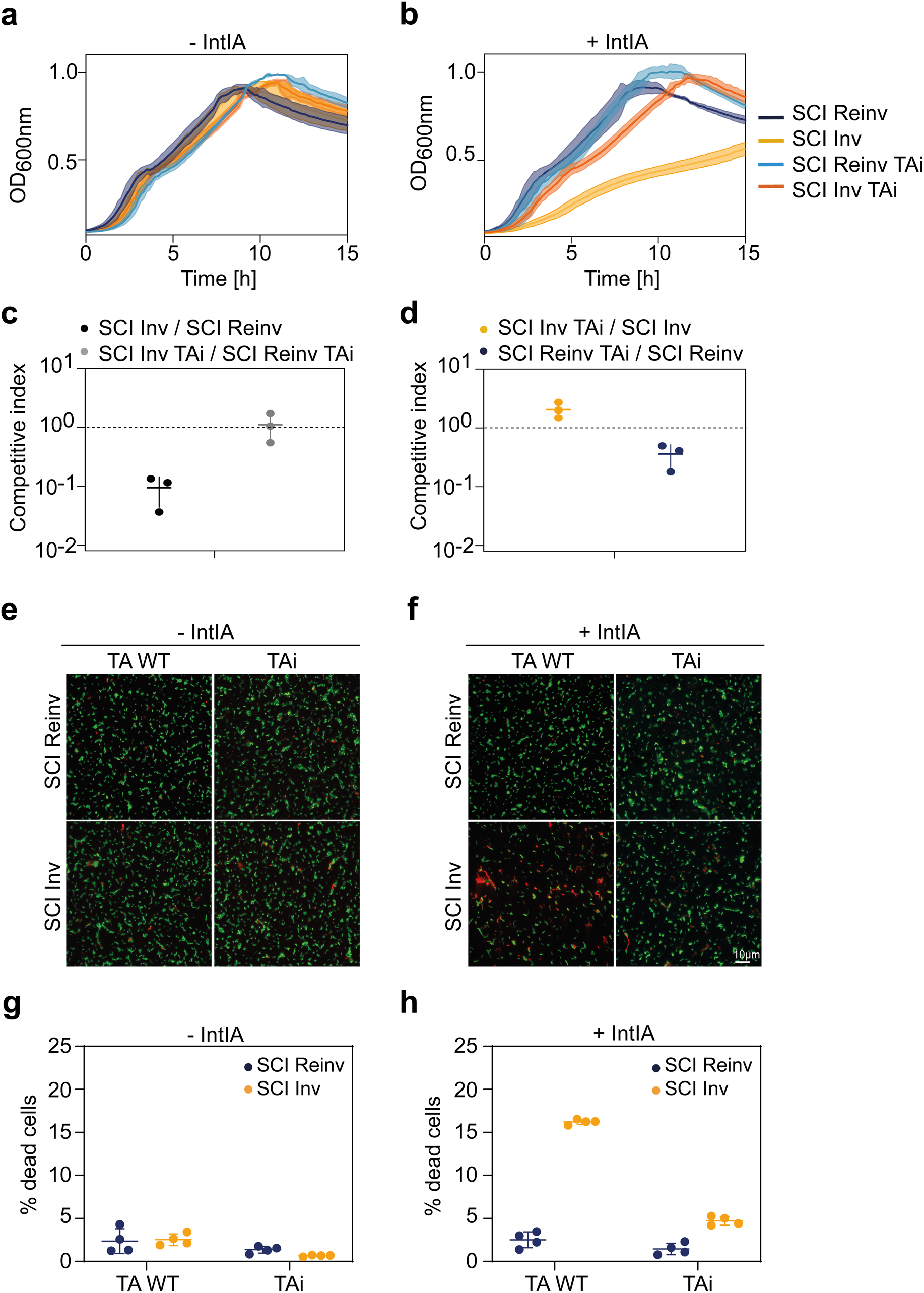
Impact of the toxin–antitoxin cassette inactivation on *V. cholerae* cell growth and viability. **a, b.** Growth curve of the strains of *V. cholerae* with its SCI Inv or Reinv and the TA WT or inactivated (TAi), in absence (a) or in presence of the IntIA integrase (b). **c.** Competitive index of the SCI Inv strain compared to the SCI Reinv strain (black) and of the SCI Inv TAi strain compared to the SCI Reinv TAi strain (grey) after 24h of co-culture. **d.** Competitive index of the SCI Inv TAi strain compared to the SCI Inv strain (orange) and of the SCI Reinv TAi strain compared to the SCI Reinv strain (blue) after 24h of co-culture. For both c and d, an index of 1 represents a ratio of 1:1 of the two strains in the mix (figured in dotted lines). **e, f.** Fluorescence microscopy images resulting from the Live and Dead assay on the SCI Inv and Reinv strains carrying TA cassettes in their WT form (TA WT) or carrying the inactivated TA cassettes (TAi), in absence (e) or in presence of the IntIA integrase (f). Green fluorescence indicates live cells (SYTO-9) whereas red fluorescence (Propidium iodide (PI)) indicates dead cells. Scale bar is 10 µm. **g, h.** Quantification of cell death as a measure of cells positively stained with PI over the total number of cells counted (∼ 1000 cells per replicate) in absence (g) or in presence of the IntIA integrase (h). Four biological replicates, means and standard deviation are represented. T: toxin, A: antitoxin

Finally, although the restoration of growth is important in absence of TA systems in the SCI Inv strain, the phenotypic rescue is not complete. We reasoned that the intense recombination activity in the SCI Inv strain in presence of integrase could also affect the replication process explaining this gap. To address this, we performed Marker Frequency Analysis (MFA), a technique that gives access to the replication pattern by deep sequencing of gDNA extracted from an exponentially growing bacterial culture (*30-32*). In absence of integrase, the MFA profile seems normal for both the SCI Inv and Reinv strains (Fig S7). However, in presence of integrase, in the SCI Inv strain, a severe drop of coverage can be observed at the genomic location corresponding to the SCI specifically. This means that there is an important replication slow-down within the SCI in this condition. Therefore, we cannot exclude a slight effect of a disruption of the replication process by the intense cassette dynamics in the SCI inverted strains in addition to the TA effect.

It was possible that the lack of constraint on the cassette array in absence of TA systems could lead to a quick and massive loss of both cassettes and the associated replication defect. For this reason, we sequenced single clones stemming from the culture of 4 SCI Inv TAi independent strains to assess their cassette content after one overnight culture in presence of the integrase. Although two clones had effectively lost 83% and 52% of their cassette content (*i.e* respectively 104 and 94 cassettes), the other two had only lost 9% and 11% of their cassette content (*i.e* respectively 16 and 20 cassettes). Thus, the increased fitness of the SCI inv TAi strain is not due to a massive loss of cassettes but rather to the absence of the detrimental effect of losing TA systems as single cassettes. We conclude that the growth defect in the SCI inverted strains is mostly due to the increased excision of TA cassettes acting as “abortive system” killing the cell. Nonetheless, we cannot completely exclude a slight effect of a disruption of the replication process by the intense cassette dynamics.

## Discussion

The recent and striking evolutionary success of the integron system to annihilate antibiotic treatment of Gram-negative pathogens, relies on the complementarity and interplay between MIs and SCIs. There is ample evidence that the large and sedentary chromosomal integrons are the source of the antibiotic resistance cassettes disseminated by the small and mobile ones (*33, 34*). The SCI arrays may carry up to more than 300 mostly silent cassettes and represent several percent of the bacterial genomes (i.e. 3% in *Vibrio cholerae*) (*5, 6*). How the carriage of hundreds of silent genes, which may become useful at one time, might be selected, remains unknown. Here, we demonstrate that TA cassettes in SCIs regulate the cassette recombination dynamics and stabilize massive SCIs.

TA systems are regularly found in mobile genetic elements and contribute to their stability. In the case of *V. cholerae*, their presence all along the SCI cassette array was proposed to avoid large scale rearrangements of the cassette array leading to the simultaneous loss of dozens of cassettes (*5, 14*). Indeed, the high prevalence of repeated sequences within SCIs might lead to massive loss of cassettes by homologous recombination in absence of regularly interspaced TA modules within the array. Strangely, all TA systems described so far in SCIs were found systematically associated with their own *attC* sites, turning them into cassettes (*5, 16, 17, 35*). This is puzzling since they all contain a promoter and they do not need to be integrated into *attI* sites to be expressed. Here, we propose an additional stabilizing activity of TA modules directly linked to their association with *attC* sites and which strikingly interconnects cassette excision frequency with cell viability (Fig 7). While the excision of most cassettes of the array is completely harmless for the cell because they do not contain a promoter and are therefore not expressed, the excision of TA cassettes immediately imposes a prohibiting fitness cost. Since TA cassettes have supposedly the same chance to be excised than any other, this means that any excision event can potentially be lethal for the cell or at least lead to growth arrest. Although it may seem surprising that TA systems are associated with recombination sites, making them easy to lose and to kill its host, we argue that this could be beneficial at the population level, analogously to their role in abortive infection (Abi) in the context of phage infection (*18*). We call this process “abortive cassette excision”, where cassette excision leads to the suicide of a fraction of the population that is proportional to the overall cassette excision rate (Fig 7). The strength of this process is that the sensor (the *attC* site) and the effector (TA system) are precisely combined into one piece (a TA cassette). This process represents an emergent property of TA systems once associated with an *attC* site, which proves to be a very effective way to ensure the stability of integron by driving the evolution of cassette excision rate towards low values. The very strong fitness cost of cassette excision imposed by the random chance of excising a TA must translate into counter-selection of any condition in which the excision rate is highly increased. Here we showed that the presence of TA cassettes heavily penalizes the inversion of the SCI of *V. cholerae* which highly increases the cassette excision rate. This makes a lot of sense with respect to integron evolution. This indeed explains our recent *in silico* analyses which reveal that most of SCIs have a specific orientation relative to the chromosome replication (*13*). It fits the observations made previously that there are many TAs in large SCIs (*14*), and the current identification of their significant and perhaps ancient over-representation in the SCI of many *Vibrio* species compared to the rest of the genome. This long co-evolution of *Vibrio* species, particularly *V. cholerae*, with their integron-encoded TA systems may have enabled SCI to have access to a vast repertoire of cassettes and thus expand their genetic capacitance. Strikingly, among the 1423 complete mobile integrons carried by plasmids, only one TA cassette was found (specifically in the MI of the 248 kb plasmid from the bacterium *Comamonas testosteroni* (GCF_014076475.1)) (*36*). This very low percentage of TA cassettes within MIs perhaps means that their recruitment can occur but lead to the abortive excision of the TA cassettes due to a too high cassette excision frequency in MIs compared to SCIs (*13*).

**Figure 7:**
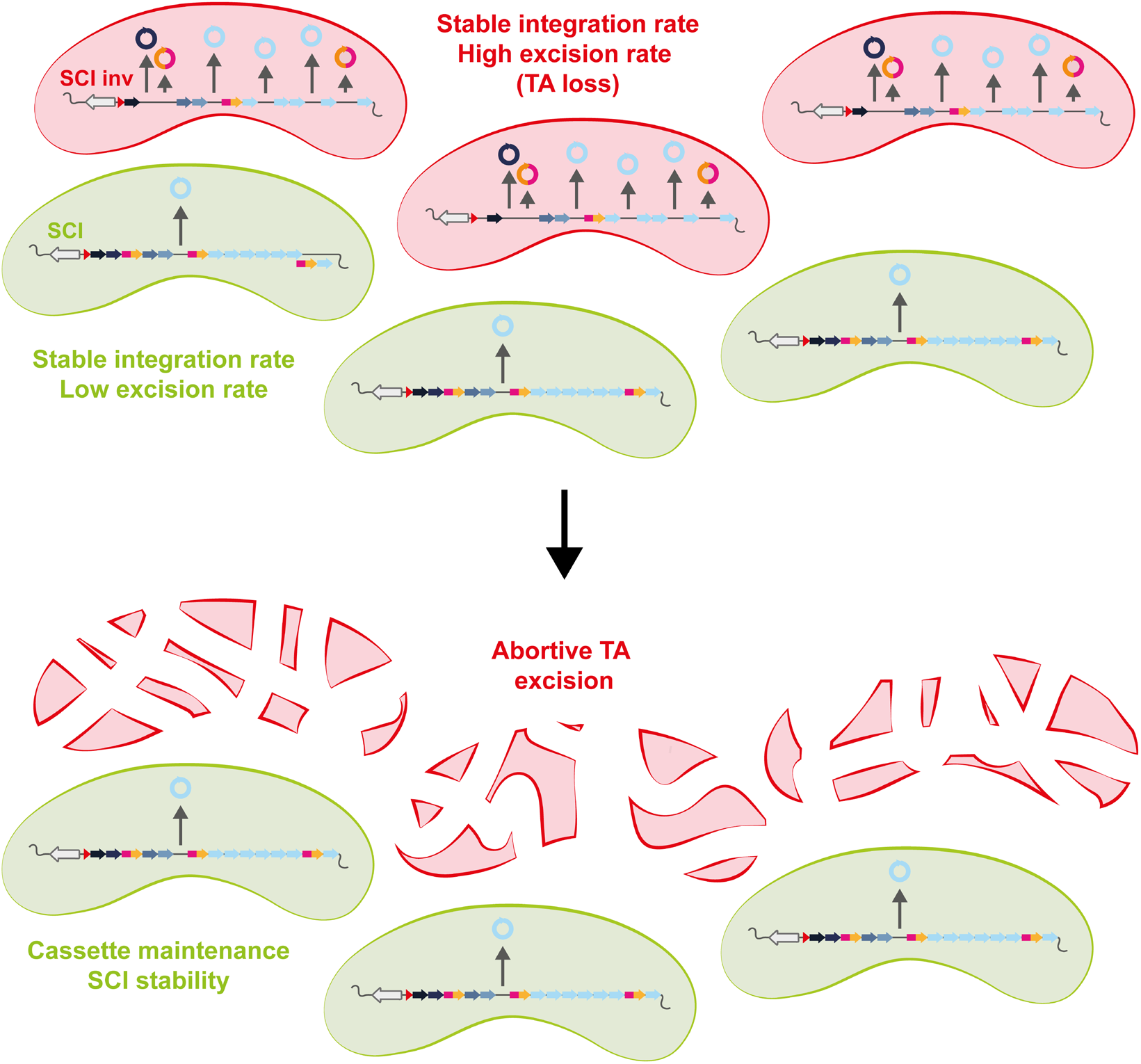
Model of the “Abortive cassette excision” process in an integron array containing toxin–antitoxin cassettes. In conditions for which the excision cassette rate is low, the TA cassettes (in pink and orange) are seldom lost. If the excision rate is high, the TA cassettes are highly excised and kill cells. This “abortive model” of TA cassette excision ensures the SCI cassette maintenance. T: toxin, A: antitoxin

Finally, in addition to their known role in plasmid stability or viral defense (*18*), TAs associated with integron recombination sites represent special cassettes largely widespread in the large SCIs and acting as selective pressure to ensure the interplay between host genome organization and genome plasticity through that we called “abortive cassette excision”.

## Materials and Methods

### Bacterial strains, plasmids and primers

The different bacterial strains, plasmids and primers that were used in this study are described respectively in Table S1, S2 and S3.

### Media

*Vibrio cholerae* and *Escherichia coli* strains were grown in Luria Bertani (LB) at 37°C. *V. cholerae* strains containing a plasmid with a thermo-sensitive origin of replication were grown at 30°C. Thymidine (dT) and diaminopimelic acid (DAP) were supplemented, when necessary, to a final concentration of 300 μM. Glucose (Glu), L-arabinose (Ara) and Fructose were added respectively at final concentrations of 10g/L, 2g/L and 10g/L. X-Gal was used at a final concentration of 100 µM. DAPG was used at a final concentration of 50 µM. Antibiotics were respectively used at the following concentrations (for *V. cholerae* and *E. coli* respectively): carbenicillin (100 μg/ml; 100 μg/ml), chloramphenicol (5 μg/ml; 25 μg/ml), kanamycin (25 μg/ml; 25 μg/ml), rifampicin (1 μg/ml; 150 μg/ml), spectinomycin (100 μg/ml or 200 μg/ml in presence of glucose; 50 μg/ml), zeocin (50 μg/ml; 50 μg/ml). To avoid catabolic repression during various tests using arabinose as inducer for the expression of the integrase, cells were grown in a synthetic rich medium: MOPS Rich supplemented with Fructose (1%) as a carbon source and arabinose (0.2%) as an inducer.

### SCI inverted and reinverted strain constructions

The inversion of the SCI was performed using a genetic tool developed in the lab (*37*) and designed to target the relocation of chromosomal DNA with bacteriophage attachment sites (HK in this case). Two DNA fragments were inserted respectively upstream and downstream of the SCI in the N16961 hapR^+^ strain (8637). The upstream fragment contained a kanamycin resistance gene and the 5’ end of a carbenicillin resistance gene associated with the *attR_HK_* site. The downstream fragment contained a zeocin resistance gene and the partner 3’ end of the carbenicillin resistance gene associated with the *attL_HK_* site (Fig S2). The strain containing the upstream and downstream fragments prior to inversion is referred to as the parental strain (Par). The expression of the HK_022_ integrase and excisionase (as a directional factor, pA401) in the parental strain led to the *attR_HK_ × attL_HK_* reaction resulting in the inversion of the SCI and to the reconstitution of the full copy of the *bla* gene, making it possible to select on carbenicillin for clones with an inverted SCI (Fig S2). As a control for the following experiments, the SCI was re-inverted to its original orientation using solely the HK_022_ integrase (p8507) to perform the *attP_HK_ × attB_HK_* reaction in the SCI inv strain.

#### Parental strain

Successive natural transformations of 8637 strain with PCR fragments produced from pF384 (fragment Kan^R^) and pF850 (fragment Zeo^R^) were performed. We used o4286 and o4302 pairwise primers to produce the Km^R^ fragment and o4657 and o4662 to produce the Zeo^R^ fragment.

#### SCI inverted strain

Transformation of I857-859 strains by the pA401 plasmid were performed at 30°C and in presence of glucose 1% to repress the HK integrase excisionase expression. Transformants were selected on spectinomycin (the marker resistance carried by the plasmid) and glucose containing plates at 30°C. The protocol of transformation is described above. SCI inversion was performed by cultivating the obtained transformant clones during 12H (overnight) at 30°C in presence of spectinomycin and arabinose 0.2 % (to induce the HK integrase excisionase expression). SCI inverted clones are selected by plating the resulting culture on plates containing carbenicillin and at 37°C (to favor the loss of the pA401 thermosensitive replication plasmid).

#### SCI reinverted strain

Transformation of the SCI inverted strain by the p8507 plasmid was transformed at 30°C (to repress the thermoinducible HK integrase promoter). Transformants were selected on Spectinomycin (the marker resistance carried by the plasmid) containing plates at 30°C. The protocol of transformation is described above. SCI re-inversion was performed by cultivating the obtained transformant clones up to OD_600_ ∼0.3 at 30°C and by shifting the temperature to 37°C during 90 min. SCI reinverted clones are selected by plating the resulting culture on plate without carbenicillin at 42°C (to favor the loss of the p8507 thermosensitive replication plasmid). The reinversion of the SCI was checked by confirming the carbenicillin sensitivity of several obtained clones (by plating them on carbenicillin containing plates).

### Automated growth curve measurements

Overnight (ON) cultures of the indicated strain were diluted 1/1,000 and then grown for 20h in the indicated medium. Bacterial preparations were distributed by triplicate in 96-well microplates. Growth-curve experiments were performed using a TECAN Infinite microplate reader, with absorbance measurements (600 nm) taken at 10-min intervals. Slopes during exponential phase were directly obtained using the “GrowthRates” R package.

### Toxin–antitoxins inactivation by allelic exchange

We performed allelic exchange to construct N16961 lacking the TA systems that could not be targeted using the Base-editing tool. To this purpose, we constructed and used different variants of the pMP7 vector, respectively pP897, pP898, pP900 and pP902. We followed the same protocols as previously described (*29*). Briefly, the suicide vector pMP7 contains a R6K origin of replication and its replication is then dependent on the presence of the Π protein in the host cell. The Π3813 cell, a *pir+* CcdB resistant *E. coli* strain, was used for cloning the different pMP7 plasmids. Once constructed, these vectors were transformed into the β3914 donor strain to deliver by conjugation the pMP7 vector into the desired recipient strain. Homology regions corresponding to the genomic DNA from the recipient strain have been cloned in the different pMP7 vector to allow the integration of the plasmid by homologous recombination. The only way for pMP7 vector to replicate into recipient strains is then to integrate into the host genome after a first crossover. After conjugation, integration of the entire pMP7 vector were then selected by plating cells on Cm plates lacking DAP. Next, cells were grown in presence of L-arabinose (0.2%) to express the CcdB toxin. The expression of this toxin allows to kill cells in which the second crossover that leads to the excision of pMP7 backbone did not take place. This method allows us to obtain mutants of *V. cholerae* that are devoid of any antibiotic resistance marker. Since we did not performed deletions, the verification of the gene replacement by PCR only was prohibited. For this reason, the STOP containing version of the toxin was also associated to a BamHI restriction site to allow an easier screening of the correct gene replacement by digesting the appropriate PCR and looking for a restriction profile. After that, the PCR products containing the BamHI sites were checked by sequencing. The primers used for PCR screening and sequencing are described Table S3.

### Golden Gate Assembly

By default, the vector expressing dCas9 carries a “random” guide that does not target any locus in *E. coli* nor in *V. cholerae* but that do contain two BsaI restriction sites (*38*). To change the guide, we perform a “Golden Gate Assembly” as such: two oligonucleotides have to be designed in the form 5’–TAGTNNNNNNNNNNNNNNNNNNNN–3’ and 5’-TTTGNNNNNNNNNNNNNNNNNNNN-3’ where the 20 “N” is the targeted sequence. For the annealing, 6uL of each oligo is mixed with 2uL of T4 DNA Ligase Buffer and 0.4 µL of T4 PNK for a total volume of 20 µL and then incubated for 30 min at 37°C. We then add 1 µL of NaCl (1M), incubate 5 min at 95°C and finally let slowly cool down to RT for at least 4h.

To insert the desired guide into the dCas9 expressing vector, we prepare the following mix: 2 µL of the dCas9 plasmid, 2 µL of annealed oligos, 1 µL of Cutsmart buffer, 1 µL of BsaI enzyme, 1 µL of ATP (1mM), 1 µL of ligase, 2 µL H_2_0. We then incubate the mix in a thermocycler with the following steps: 3 min at 37°C (for digestion), 4 min at 16°C (for ligation) and alternate between those steps 25 times. We finish with one cycle of 5 min at 50°C and 5 min at 80°C for enzymes inactivation. At this step, the mix is ready to be transformed in a cloning *E. coli* strain and the successful sgRNA insertions are screened by PCR using the o2406 and o3812 primers.

### Base editing

The tool used for base editing is derived from the catalytically dead Cas9 (dCas9) previously described (*27*). The construction consists in a gene fusion between dCas9 and CDA1, a cytidine deaminase that catalyzes the deamination of cytidine, resulting in a uridine base which will later be replicated as a thymine (C → U→ T). A linker of 100 AA separated the dCas9 and the CDA1. An Uracil Glycosylase Inhibitor (UGI) was also fused to CDA1 to increase the base editing rate as described by Banno and coll as well as a LVA degradation tag to decrease the toxicity the construct (*27*). Due to a low efficiency transformation rate of this construct in *Vibrio cholerae*, the construct was delivered by conjugation. Hence, the appropriate construct was first transformed in a β3914 donor strain. Then both the donor and receptor strains were cultivated to exponential phase, mixed 1:1 in a 2 mL tube and centrifugated (6000 rpm for 6 min).

The pellet was spread on a membrane on a plate containing DAP to sustain growth of the donor strain and DAPG for induction of base editing, and the plate was incubated for 3h at 37°C. After incubation, the membrane was resuspended in 5mL LB and vortexed for 30s to resuspend the conjugated cells which were then plated on appropriate media: Cm_5_ + Glc1% + Xgal to obtain both the CFU/mL after conjugation and the base editing efficiency or Rif_1_ + Cm_5_ + Glc1% to obtain the rate of apparition rifampicin resistant clones as a proxy for global mutation rate. Blue colonies had a functional *lacZ* gene while the white colonies were synonym for a mutated *lacZ* gene. All targeted sites were checked by PCR and sequencing (see Table S3). Then the whole genomes of the TAi strains were sequenced (Illumina).

### SCI cassette shuffling assay

A single clone of each strain was isolated on plate and then inoculated for an overnight culture in LB + Spec + Glc 1%. The next day, the cells were inoculated at 1/50^th^ for 2h in MOPS Rich + Spec + Fructose 1% + Arabinose 0.2% as a pre-induction step until they were at exponential phase. They were then inoculated at 1/1000^th^ and grew for 20h until stat phase. Cultures were streaked on 1% agarose plates containing MH+ Spec + Glc 1% and single clones were isolated. A total of 48 PCR per condition was performed using an oligo (o5778, see Table S3) hybridizing in the promoter of *intIA* (stable part) and one (o1401) hybridizing in the first cassette of the array (variable part). A representative sample of the PCR profile for each condition was migrated in 1% agarose for visualization.

### Suicide conjugation assay

This assay has been previously described by Vit and coll. 2021 (*3*). We used the suicide conjugative vector pSW23T (*39*) that allows the delivery of the bottom strand of the *attC_aadA7_* recombination site (pD060). For each condition assay, at least 16 recombinant clones were isolated on appropriate plates and analyzed by PCR. To determine precisely if the pSW23T vector has been inserted into the *attIA* site of the SCI we used 5778 and SWend primers. These primers hybridize respectively in a sequence upstream of *attIA* in *V. cholerae* chromosome 2 or downstream of *attC_aadA7_* in the pSW23T vector. For each condition assay, at least three PCR reactions were purified and sequenced to confirm the insertion point.

### SCI cassette excision assay

A single clone of each strain was isolated on plate and then inoculated for an overnight culture in LB + Spec + Glc 1%. The next day, the cells were inoculated at 1/50^th^ for 2h in MOPS Rich + Spec + Fructose 1% + Arabinose 0.2% as a pre-induction step until they were at exponential phase. They were then inoculated at 1/1000^th^ and grew for 20h until stat phase. Finally, 1mL of this culture was used to inoculate a 100 mL culture in LB + Spec + Glc1 % to filter the dead cells of the latter culture. The resulting culture constitutes a mixed population where each individual might have experienced a diverse set of cassette excision events. Bulk DNA was extracted from these mixed population (Qiagen genomic DNA extraction kit) and sequenced using the Pacific Bioscience (PacBio) long-read sequencing technology. Mapping was performed using minimap2 (*40*) and the cassette deletions were detected using a homemade R program. In order not to confuse the excision rate with the random chance of an excision event to be propagated, the cultures of the different strains were replicated eight times independently.

The cassette excision frequency corresponds to the total number of cassette excision events divided by the sequencing depth across the SCI. For the SCI inverted strain expressing integrase, we detected 478 excision events encompassing the entire SCI. The calculated frequency corresponds to: N_exc_ / (SCI_cov_ + N_exc_); where N_exc_ is the number of excision events and SCI_cov_ is the mean coverage for each cassette multiplied by the number of cassettes in the SCI (i.e. 179). We therefore obtained a cassette excision frequency of 478 / (13269.16 + 478) = 3.48e-02.

For the SCI inverted strain without integrase and the SCI reinverted strains with and without integrase, for which we did not detect any excision event, we calculated the limit of detection. It corresponds to: 1/ SCI_cov_. Note that the SCI read coverage obtained for each condition is respectively 13518.08, 14737.07 and 14550.91 corresponding to detection limits of respectively 7.40e-05, 6.79e-05 and 6.87e-05.

### Protein purification

IntIA proteins were expressed in BL21 bacteria (*41*). Cells were lysed in 25 mM Tris pH 8, 1M NaCl, 1 mM PMSF, lysozyme and protease inhibitors containing buffer. Proteins were purified by nickel affinity chromatography with a 200 mM imidazole elution. His-tag was removed by incubating proteins at 4°C overnight with TEV protease (1 mg per 15 mg of total proteins containing a 5 mM final concentration of β mercapto-ethanol). After dilution to 150 mM NaCl, proteins were purified on an SP column equilibrated with 25 mM Tris pH8 and 150 mM NaCl. A gradient of 150 mM to 1M NaCl was performed. Proteins were then brought to a final concentration of 1 mM DTT and 10% glycerol.

### Electromobility shift assay

Each reaction contained 500 ng Poly[d(I-C)], 12 mM Hepes–NaOH pH 7.7, 12% glycerol, 4mM Tris–HCl pH 8.0, 60 mM KCl, 1 mM EDTA, 0.06 mg/ml BSA, 1 mM DTT, 10% Tween20, 0.01 nM specified 32P-labeled DNA oligonucleotide (Table S3) and the specified quantities of purified His-tagged IntIA and IntIAY302F, in a final volume of 20 μl. The samples were incubated at 30 °C for 10 min without the probe followed by 20 min with the probe, then loaded to a 5% native polyacrylamide gels with 0.5x TBE as running buffer. The gels were visualized using a Molecular Dynamics intensification screen and a Typhoon FLA 9500 laser scanner

### Competition assay

The SCI Inv strains are resistant to Carb, which is not the case for the SCI Reinv strains (see above). We used this difference to perform a competition assay. One clone of each strain was isolated on plate and used to inoculate an overnight culture. Before to start the competition assay, the cells were precultured at 1/50^th^ in LB + Spec + Glc 1% until exponential phase to be not biased by the dead cells from the overnight culture. Cells were then mixed at 1:1 ratio at 1/200^th^ in MOPS Rich + Fructose 1% + Spec + Arabinose 0.2% for induction of the integrase. The co-culture was plated on petri dishes containing MH + Spec + Glc 1% to get the total CFU/mL and in parallel on MH + Carb + Spec + Glc 1% plates to get the number of SCI Inv CFU/mL in that same culture. We plated at t_0_, t_0_ + 2h, t_0_ + 4h, t_0_ + 6h, t_0_ + 8h, t_0_ + 24h. The competitive index (I) was calculated as such: I = CFU_SCI Inv_ / CFU_SCI Reinv_ – CFU_SCI Inv_ so that an index of 1 represents a situation where the ratio of SCI Inv and SCI Reinv is 1:1 (Neutrality). A high index indicates that SCI Inv has a competitive advantage compare to its Reinv counterpart, and *vice-versa*.

### Live and dead assay

Overnight cultures were performed in LB + Spec + Glucose 1%. Then day cultures were inoculated at 1:1000 inMOPS Rich + Spec + Fructose 1% + Arabinose 0.2% medium to OD600nm⁓ 0.8. Then 50 µL of the latter culture was dyed using the LIVE/DEAD™ *Bac*Light™ Bacterial Viability Kit, that is a mix of Propidium Iodide (PI) and SYTO-9 dyes. PI specifically stains the dead cells in red. SYTO-9 specifically stains viable cells in green. Cell viability was then assessed by microscopy. Stained cells were placed of 1.4% agarose pad containing MOPS Rich and observed using a Zeiss ApoTome inverted wide-field microscope. Snapshot images were taken with a 40-ms and 50-ms exposure time (FITC and DsRED channel respectively), using a Plan Apo 63× objective (numerical aperture = 1.4, +optovar 1.6×) using a Hamamatsu sCMOS ORCA-Flash 4.0 v3 (Institut Pasteur imaging facility imagopole). The result is presented as the proportion of dead cells (red) in the population.

### Microscopy setup for live imaging

Bacterial cells were grown to mid-exponential phase in liquid MOPS Rich + Spec + Fructose 1% + Arabinose 0.2% medium and then transferred to 1.4% agarose-padded slides containing MOPS Rich + Spec + Fructose 1% + Arabinose 0.2%. A coverslip was placed on top of the agarose pad and sealed with a vaseline:lanolin:paraffin mix (ratio 1:1:1) to prevent pad evaporation. Slides were incubated at 37°C under an inverted wide-field microscope (Zeiss ApoTome) for time-lapse video recording. Video frames were taken at 1 min interval time for a total duration of 170 min, using a Plan Apo 63× objective (numerical aperture = 1.4) using a Hamamatsu sCMOS ORCA-Flash 4.0 v3 and autofocus module (Definite focus, Zeiss) (Pasteur Institute Imaging Facility Imagopole). A total of 8 movies were recorded and analyzed across all strains.

### Marker Frequency Analysis

MFA was performed as described in (*32*). Briefly, cells were grown in MOPS + Spec + Arabinose 0.2% + Fructose 1% at 37°C under agitation. Genomic DNA was prepared using the DNeasy® Tissue Kit (Qiagen) from 30ml of exponentially growing cells (OD_450_ ∼0.15). The remaining culture was washed twice and was kept under agitation in MOPS + Spec + Glucose 1% at 37°C to repress expression of IntIA and to prepare gDNA from cells in stationary phase (24h of culture). Libraries were prepared for Illumina sequencing (150 bp, paired end). The resulting reads were mapped with bowtie2 (*42*). The genomic coverage of the sequencing data stemming from the exponentially growing cultures of each strain was normalized with the corresponding data in stationary phase. The normalized coverage is represented in function of the relative position to the origin of replication of each chromosome as colored dots for 1 kb bins and red dots for 10 kb bins. Dots are not shown when at least one position has a coverage of zero within the bin (typically due to repetitive sequences that prevents correct reads mapping). The gradient of coverage from *ori* to *ter* serves as a proxy for the replication dynamics and the slope translates the local speed of the replication fork (*31*).

### Comparative genomics analysis of toxin–antitoxin systems within SCI

#### Genome dataset and SCI annotation

To study the enrichment of TA systems within SCI across bacterial genomes, we used the dataset and integron predictions of IntegronFinder 2.0 (*20*). Briefly, this dataset consists of 21,105 complete genomes retrieved from NCBI RefSeq on 30 March 2021. We defined SCIs as complete integrons with at least 11 predicted *attC* sites. Several isolates of *Vibrio cholerae* appeared to carry large (>10 *attC* sites) CALINs (cluster of *attC* sites lacking an integrase) (*20, 43*). A closer investigation showed that these CALINs were preceded by a pseudogenized integrase, which relates them with SCIs. These elements could have been recently inactivated, or the putative pseudogenes could result from sequencing artifacts. Hence, we included in our analysis all the isolates carrying these CALINs. To simplify the results, we grouped together SCIs and CALINs, without distinction, under the global term SCIs.

#### Toxin and antitoxin prediction

In each genome harboring at least one SCI, we predicted toxins and antitoxins with TASmania, a predictor with very low false negative rate (*21*). Our goal was to retrieve even unusual TA systems that would be rarely found outside SCIs and hence missed by more conservative predictors. More precisely, we downloaded toxin and antitoxin HMM models from TASmania’s website (accessed on 25 November 2021) and screened for them with the hmmscan command of HMMER 3.3 (*44*) (Nov 2019; http://hmmer.org/). As the command was run with default parameters, no threshold was used to filter hits in the first screening. To reduce the number of false positives, we evaluated TASmania’s performance on the genome of *Vibrio cholerae* N16961’s secondary chromosome, known to harbor 19 TA systems (*16, 17*), all located within the SCI. We observed that applying an e-value threshold of 10^-3^ allowed to keep all the hits corresponding to characterized TA systems (as described by Iqbal and coll. (*16*)), while eliminating most of the other hits. Hence, we selected this value to filter TASmania hits in all the genomes comprised in our analysis and we decided to consider as a toxin (respectively antitoxin) hit any protein matching a TASmania toxin (respectively antitoxin) profile with an e-value lower than 10^-3^.

#### Contingency analysis of TA enrichment within SCI

For each genome, we performed a contingency table analysis to test for an enrichment of TA systems within SCIs. The contingency table was built as following: in each genome, each protein was classified as, on one side, belonging or not to an SCI, and, on the other side, containing a TASmania hit or none. A Fisher one-sided test for statistical enrichment significance was performed on each contingency table. To correct for multiple testing, the resulting p-values were adjusted either with the Benjamini-Hochberg (main analysis) or the Bonferroni (complementary analysis) corrections. An isolate was claimed to harbor a SCI significantly enriched in TA when the adjusted p-value was lower than 0.05. The Benjamini-Hochberg and Bonferroni corrections were re-computed in the analysis focusing on *Vibrio* isolates.

#### Vibrio genus cladogram

To identify a potential evolutionary trend explaining the TA enrichment within SCI across the *Vibrio* genus, we reconstructed a cladogram grouping *Vibrio* species together. More precisely, we took as basis the phylogeny published by Sawabe and coll. (*22*) and removed the species that did not harbor any SCI. The species comprised in our dataset that were absent of the phylogeny were then added as a sister branch of the closest relative species.

## Acknowledgments

We thank Samuel H. Sternberg for critical reading of the manuscript. We thank Jun Teramoto and Akihiko Kondo for sharing their construct for base-editing. We thank Marc Monot, Laurence Ma, Juliana Pipoli Da Fonseca and Thomas Cokelaer from the Biomics platform, C2RT, Institut Pasteur, Paris, France, supported by France Génomique (ANR-10-INBS-09) and IBISA. We thank Clarisabel Garcia Rodriguez for her help with microscopy.

## Funding

Institut Pasteur

Centre National de la Recherche Scientifique (CNRS-UMR 3525)

Fondation pour la Recherche Médicale (FRM Grant No. EQU202103012569)

ANR Chromintevol (ANR-21-CE12-0002-01)

French Government’s Investissement d’Avenir program Laboratoire d’Excellence ‘Integrative Biology of Emerging Infectious Diseases’ [ANR-10-LABX-62-IBEID]

Ministère de l’Enseignement Supérieur et de la Recherche and the Direction Générale de l’Armement (DGA)

## Author contributions

Conceptualization: ER, CL, DM, DB, EPCR

Methodology: ER, CL

Investigation: ER, BD, EL, CV, CW, JB, FF, GG, VC, DL, VP, FR, OS, CL

Supervision: CL, DM, DB, EPCR

Writing—original draft: ER, CL, DM

Writing—review & editing:

## Data and materials availability

All data are available in the main text or the supplementary materials. MFA and PacBio sequences are publicly available in The European Nucleotide Archive (ENA). For the MFA analysis, the accession no. are ERR11475429, ERR11475428, ERR11475427, ERR11475426, ERR11475425, ERR11475424, ERR11475423 and ERR11475422. For the PacBio sequencing, the accession no. are ERR11475405, ERR11475404, ERR11475403 and ERR11475402. All other raw data are available upon request to the corresponding author.

