## Supplementary material for "Cassette recombination dynamics within chromosomal integrons are regulated by toxin–antitoxin systems" (from another version of this preprint: version 4, 2023-06-22)

### Supplementary figures

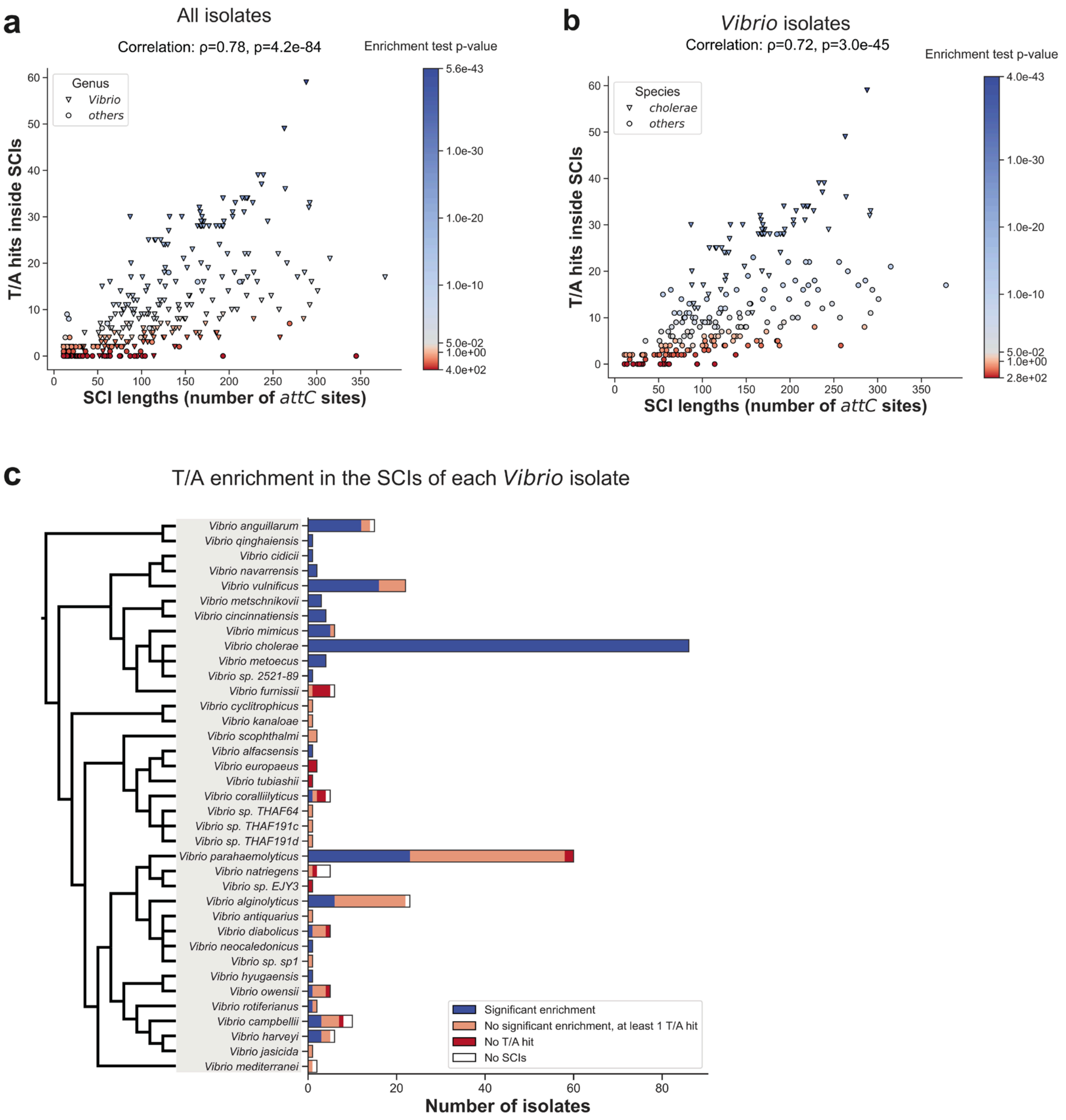


**Figure S1: Toxin–antitoxin landscape in SCIs, re-analyzed with a harsher correction.**

This figure is a re-analysis of Figure 2, applying the very conservative Bonferroni correction instead of the Benjamini-Hochberg correction.

**a, b.** Each point represents a single NCBI RefSeq isolate harboring at least one SCI (>10 *attC* sites). The x-axis displays the total number of *attC* sites in the (potentially multiple) SCI of the isolate. The y-axis exhibits the total number of TASmania toxin and antitoxin HMM hits in the SCI of the isolate (e-value<$10^{-3}$). The Spearman nonparametric rank correlation coefficient is

shown at the top of the graph, together with its significance p-value. Point hue indicates the Bonferroni adjusted p-value of Fisher's test for TASmania hits enrichment in SCI compared to the rest of the genome (scale on the right of the graph, significant enrichment in blue). **a.** All the isolates harboring at least one SCI are displayed. The shape of each point on the graph represents whether the isolate belongs to the *Vibrio* genus (triangles) according to the NCBI taxonomy or to another genus (circles). **b.** Same graph as a, but only with isolates belonging to the *Vibrio* genus. The shape of each point on the graph represents whether the isolate belongs to the *Vibrio cholerae* species (triangle) or to another species (round).
**c.** Distribution of *Vibrio* isolates by species according to the toxin/antitoxin enrichment inside the SCI they harbor.
T: toxin, A: antitoxin

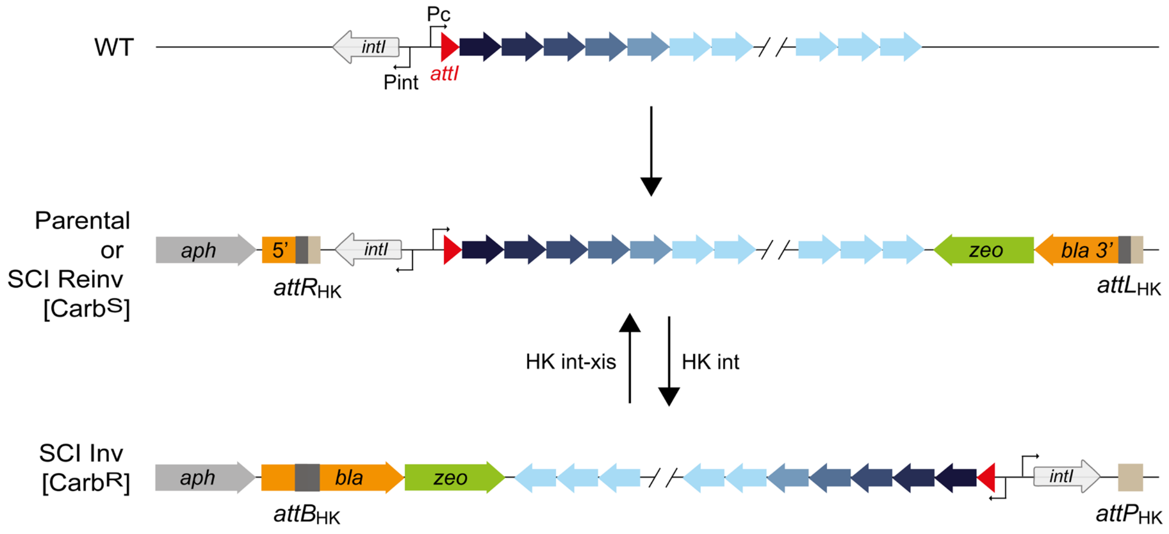


**Figure S2: Construction of the *V. cholerae* strains with the SCI inverted and reinverted.**

Schematic of the process of inversion/reinversion of the SCI in *V. cholerae*. The SCI was flanked by two fragments containing the HK phage recombination sites. Upstream, the *aph* gene conferring resistance to kanamycin was inserted alongside the $attR_{HK}$ site and the 5' end of the *bla* gene conferring resistance to carbenicillin ($Carb^R$). Downstream, the *zeo* gene conferring resistance to zeocin was introduced alongside the $attL_{HK}$ site and the 3' end of the *bla* gene. Expression of both the HK phage integrase and excisionase (HK int-xis) induced the inversion of the SCI and the reconstitution of the *bla* gene, giving $Carb^R$ SCI Inv clones. Expression of the HK phage integrase (HK int) in those new clones induced the very efficient reinversion of the SCI, giving SCI Reinv clones ($Carb^S$).

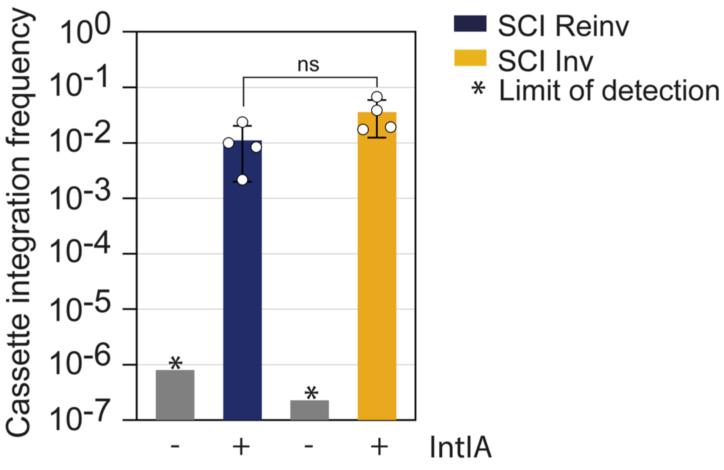


**Figure S3: Cassette integration frequencies in the *V. cholerae* SCI.**

Insertion of a synthetic cassette delivered by conjugation in *V. cholerae* with its SCI Inverted (Inv, orange bar) or Reinverted (Reinv, blue bar). The grey bars represent the frequencies obtained in absence of integrase (4 biological replicates). Asterisk (*) indicates the recombination frequency was below detection level, indicated by the bar height (limit of detection). Error bars show the sd. Statistical comparison (t test) was performed. ns. (not significant).

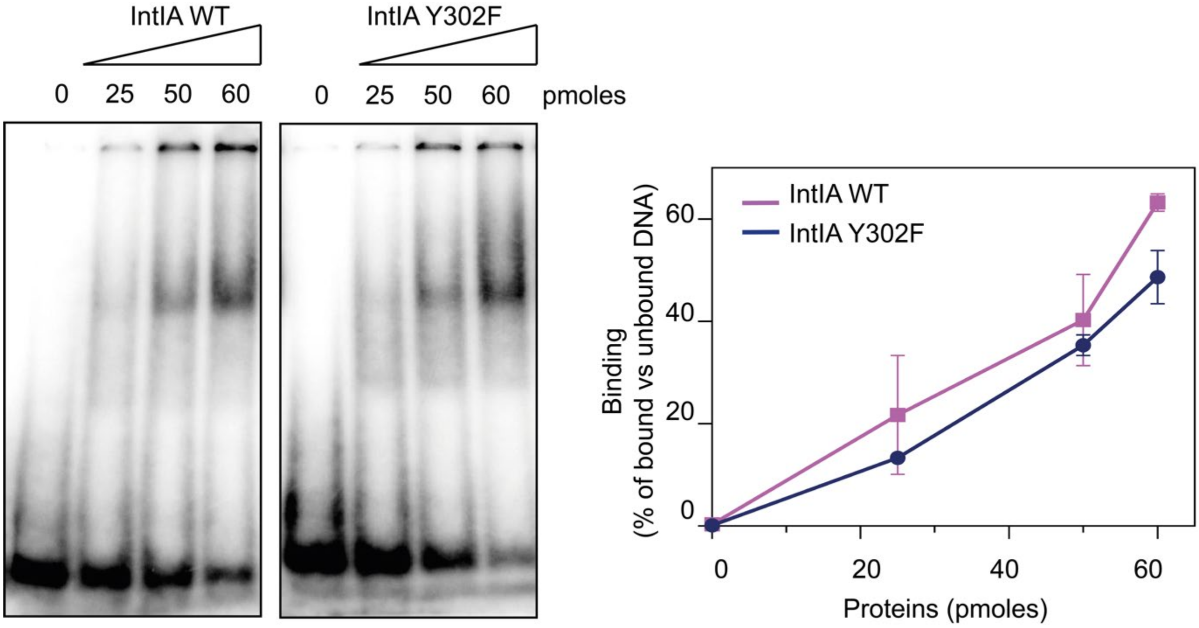


**Figure S4: Effect of loss of catalytic activity of IntIA on its *attC*bs binding capacity.**

Gel retardation assay (EMSA) was done as described in Materials and Methods in the presence of the indicated amounts of the wild type IntIA (WT) or the catalytically inactive IntIA (Y302F) and an oligomer comprising the sequence of a VCRbs. The graph on the right summarizes the outcome of 3 independent EMSA.

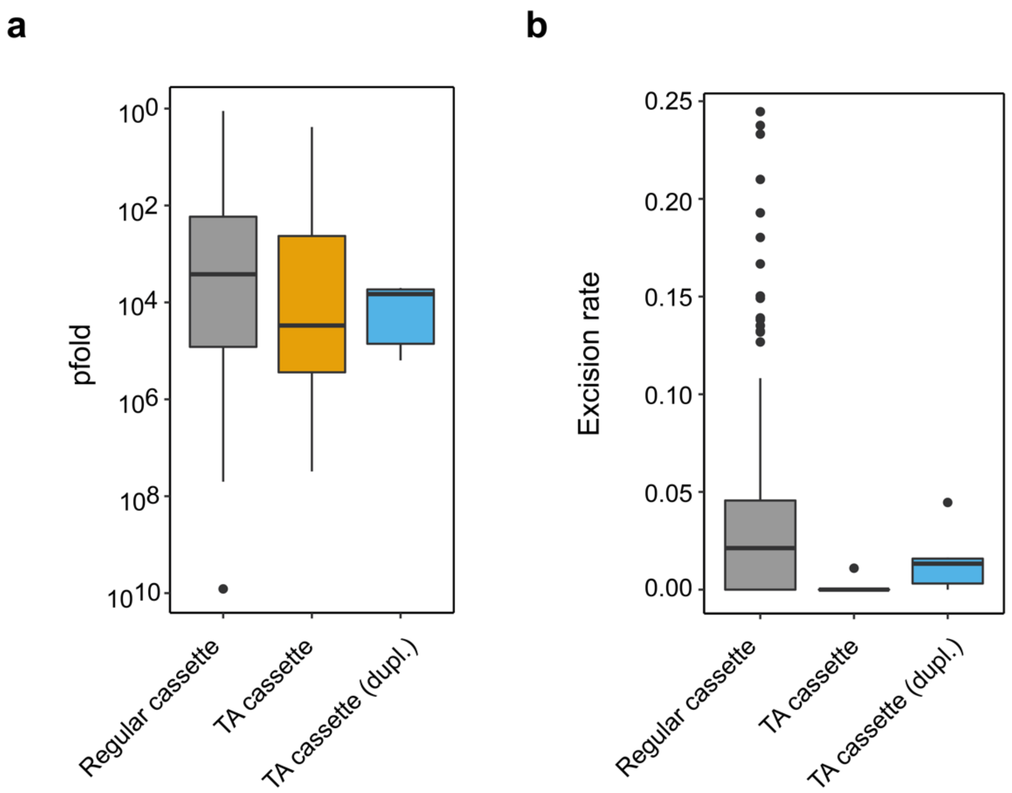


**Figure S5: Comparison of the pfold and excision rates of the regular and toxin–antitoxin cassettes in the *V. cholerae* SCI.**

**a,b.** Boxplot summarizing the distribution of respectively the pfold and excision rate values of the cassettes that do not encode a TA system (grey, left), of the cassettes that encode a TA system that is not duplicated in the array (orange, middle) and of the cassettes that encode a TA system that is found in more than one copy in the SCI. The median value is represented as a bold black line for each box. pfold was calculated as indicated in the Materials and Methods section and reflects the ability of an *attC* site to form a recombinogenic secondary structure.
T: toxin, A: antitoxin

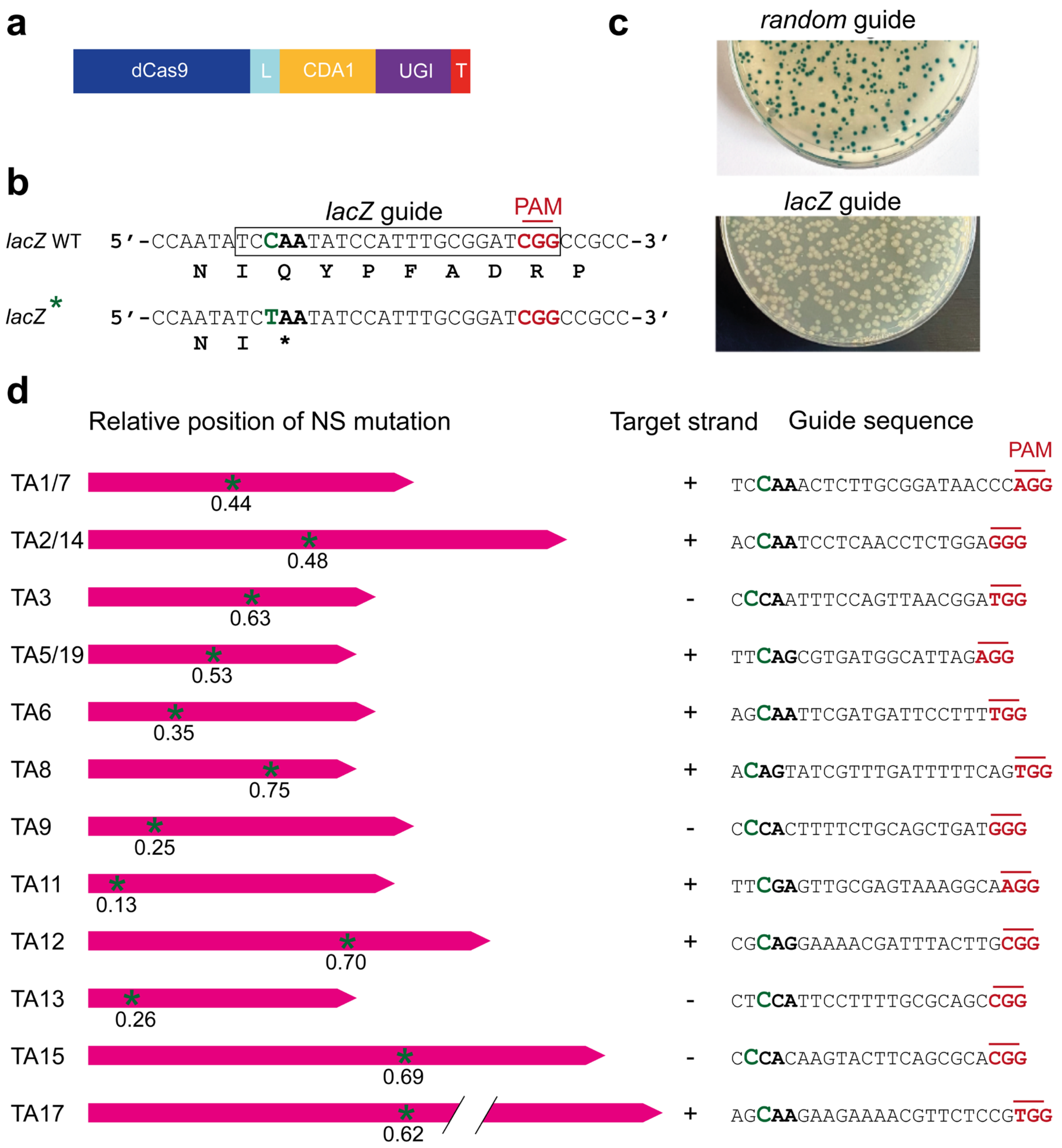


**Figure S6: Inactivation of toxin–antitoxin systems in the *V. cholerae* SCI.**

**a.** Schematic view of the construct used for base editing. From 5' to 3': dCas9, a 121 aa linker (L), the *Petromyzon marinus* cytosine deaminase (CDA1), uracil glycosylase inhibitor (UGI) and a protein degradation tag LVA (T).
**b.** Representation of the target within the sequence of lacZ used to test the efficiency of the Base-Editor with the corresponding protein sequence. The blue box represents the guide sequence and the PAM sequence highlighted in red. The modified cytosine (in bold) and the resulting NS mutation are represented below.

**c.** Xgal-containing plates of *V. cholerae* after the induction of the Base Editor directed by a random guide (up) or the *lacZ* guide (below). The white cells carry a mutation in *lacZ*, demonstrating the high efficiency of base editing in *V. cholerae*.
**d.** Detail of the targets of the Base Editor, with a representation of the targeted genes. The target is represented by a white star and the relative size of the resulting truncated protein compared to the WT version is indicated. Details on the guides are also represented.

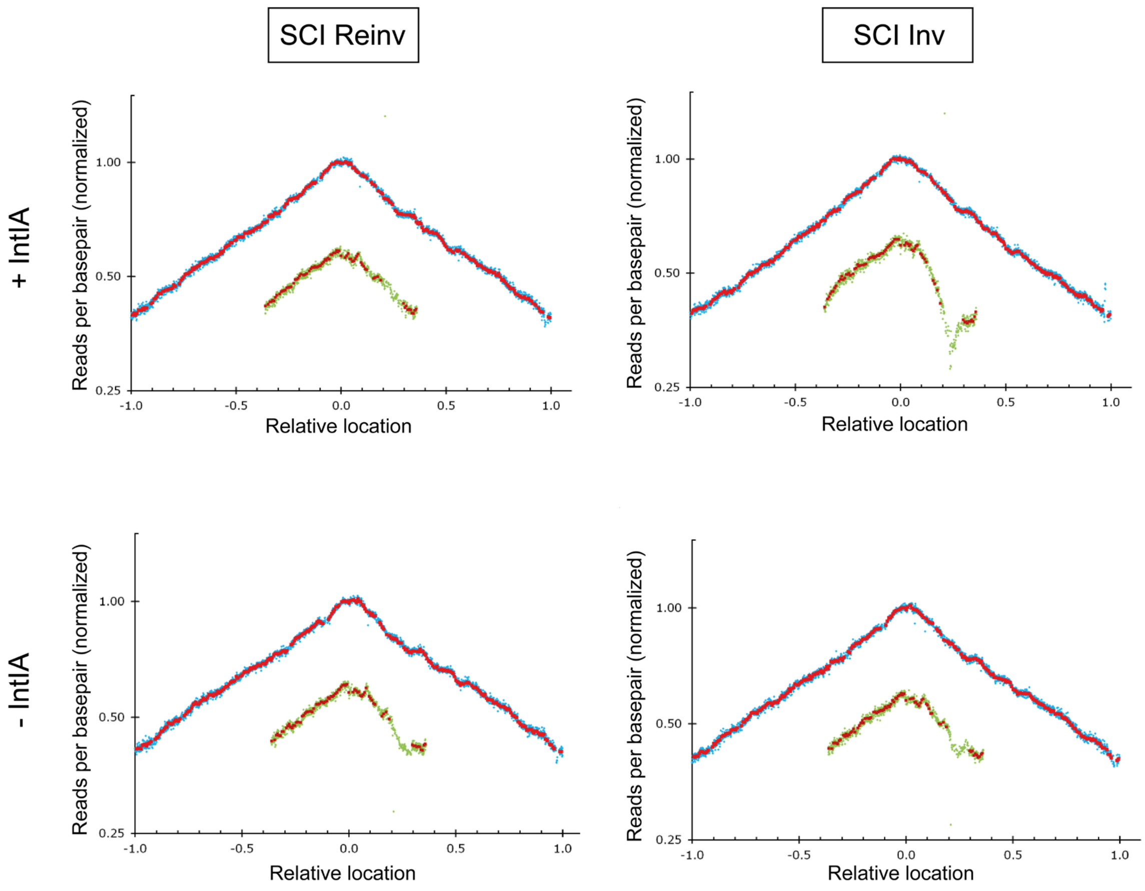


**Figure S7: Impairment of replication in the *V. cholerae* SCI Inv strain expressing the integrase.**

Marker Frequency Analysis profiles of the SCI Inv and Reinv strains expressing or not the integrase. The Y axis represents the number of reads per basepair, normalized by the value obtained for the *ori1*. In blue, the values obtained for *chr1* and in green for the *chr2*, with bins of 1kb. In red are the values computed with 10 kb bins. The graphs are centered around the *ori1* and the Y axis represents the normalized distances on the left (negative values) or the right (positive values) replichores as to compare with the size of the *chr1*. A drop of the replication speed within the SI is evidenced in the SCI Inv + IntIA by a drop of the relative coverage in that region (*chr2*, right replichore, proximal to *ter2*).

#### Supplementary tables

##### Table S1: Bacterial strains used in this study

| Strain number | Relevant genotypes or description | References |
|---|---|---|
| **Bacterial strains** | | |
| One Shot™TOP10 | Top10 (F-) mcrA Δ( mrr-hsdRMS-mcrBC) Φ80lacZΔM15 ΔlacX74 recA1 araD139 Δ(ara leu) 7697 galU galK rpsLendA1 nupG [StrR] | Thermo Fischer Scientific (Ref: C404010) |
| BL21(DE3) | $F^{-}$*ompT* *hsd*$S_B$ ($r_B^{-}$, $m_B^{-}$) *gal* *dcm* (DE3) | Thermo Fischer Scientific (Ref: C600003) |
| DH5α | $F^{-}$ φ80*lac*ZΔM15 Δ(*lac*ZYA-*arg*F)U169 *rec*A1 *end*A1 *hsd*R17($r_K$, $m_K^{+}$) *pho*A *sup*E44 $\lambda^{-}$*thi*-1 gyrA96 relA1 | Laboratory collection |
| 8637 | *V. cholerae* N16961 biovar El Tor, hapR+ [$Strep^R$] (WT) | Laboratory collection |
| L041 | 8637 Δ*attIA* | This study, *attIA* site was deleted by allelic exchange by delivering the suicide conjugative plasmid pK590 in the 8637 strain. |
| 8725 | π3813 lacIq thi-1 supE44 endA1 recA1 hsdR17 gyrA462 zei-298::Tn10 ΔthyA::(*erm-pir116*) [ErmR] | *(29)* |
| 4196 | β2163 (F-) RP4-2-Tc::Mu ΔdapA::(erm-pir) [KmR ErmR] | *(39)* |
| 8726 | β3914, β2163 gyrA462 zei-298::Tn10 | *(29)* |
| I857, I858, K022, K023, K024 | 8637 rplt-aph-bla5'-attRHK-intIA // VCA0508-ble-bla3'-attLHK-VCA0510 (parental strains) | This study |
| I992, I993, J341, J344, J350 | SCI inverted. *attCbs* on lagging strand template (SCI Inv strains) | This study |
| J053, J054, K025, K028, K030 | SCI re-inverted into its original orientation, *attCbs* on leading strand template (SCI Reinv strains) | This study |
| L042, L216, L217 | I857-I858-K022 (parental) Δ*attIA* | This study |
| L043, L218, L219 | I992-I993-J341 (SCI Inv) Δ*attIA* | This study |

| | | |
|---|---|---|
| L044, L220, L221 | J053-J054-K025 (SCI Reinv) Δ*attIA* | This study |
| R550, R556, R561 | SCI Reinv TAi | This study |
| R555, R560 | SCI Inv TAi | This study |
| S507, S513 | SCI Inv TAi, obtained from inversion of R550 and R556 respectively | This study |
| S764-S767 | SCI Inv TAi strains cultivated for 24h in presence of IntIA and isolated for gDNA sequencing | This study |
| **Transformed strains** | | |
| L120 | β2163 + pD060 | *(3)* |
| P998 | β3914 + pP897 | This study |
| P999 | β3914 + pP898 | This study |
| Q002 | β3914 + pP900 | This study |
| Q004 | β3914 + pP902 | This study |
| P388 | β3914 + pP111::$lacZ_{Vch}$ | This study |
| K591 | β3914 + pK590 | This study |
| K471 | 8637 + pJ504 | This study |
| K483, K487, K491 | SCI Inv + pJ504 | This study |
| K493, K497, K501 | SCI Reinv + pJ504 | This study |
| K503 | 8637 + pJ502 | This study |
| K515, K519, K523 | SCI Inv + pJ502 | This study |
| K525, K529, K533 | SCI Reinv + pJ502 | This study |
| K876 | 8637 + pK752 | This study |
| K888, K892, K896 | SCI Inv + pK752 | This study |
| K898, K902, K906 | SCI Reinv + pK752 | This study |
| L042, L364, L365 | SCI InvΔ*attIA* + pJ504 | This study |
| L041, L358, L359 | SCI InvΔ*attIA* + pJ502 | This study |
| L043, L366, L367 | SCI ReinvΔ*attIA* + pJ504 | This study |
| L044, L360, L361 | SCI ReinvΔ*attIA* + pJ502 | This study |
| S688-S690 | SCI Inv TAi + pJ504 | This study |
| R827, R830, R835 | SCI Reinv TAi + pJ504 | This study |
| S685-S687 | SCI Inv TAi + pJ502 | This study |
| R848, R851, R856 | SCI Reinv TAi + pJ502 | This study |

**Table S2: Plasmids used in this study**

| Plasmid number | Plasmid description | Relevant properties and construction |
|---|---|---|
| p979 | pBAD18 | *ori*ColE1; [Carb$^R$] *(3)* |
| pA401 | pSC101*rep*$^{TS}$:: *int$_{HK}$-xis$_{HK}$* | *oripSC101rep*$^{TS}$ *oriT*$_{RP4}$; [Sp$^R$] (Bland, unpublished) |
| p8507 | pSC101*rep*$^{TS}$:: *int$_{HK}$* | *oripSC101rep*$^{TS}$ *oriT*$_{RP4}$; [Sp$^R$] (Bland, unpublished) |
| pF384 | pTOPO*::rplT-aph-bla*5'-*att*RHK-*intIA* | *ori*ColE1; [Km$^R$] (Vit, unpublished) |
| pF850 | pTOPO*::VCA0508-ble-bla*3'-*att*LHK-*VCA0510* | *ori*ColE1; [Km$^R$] (Vit, unpublished) |
| pG086 | pBAD18*::arr2* | *oriColE1*; [Rif$^R$]<br>*arr2* cloned into PsiI/SphI digested p979. |
| pJ502 | pBAD43*ΔattC$_{aadA1}$* | *oripSC101*; [Sp$^R$] *(3)* |
| pJ504 | pBAD43 *ΔattC$_{aadA1}$::intIA$_{Vch}$* | *oripSC101*; [Sp$^R$] *(3)* |
| pK752 | pBAD43 *ΔattC$_{aadA1}$::intIA$_{Vch}$* (Y302F) | *oripSC101*; [Sp$^R$]<br>Quick change mutagenesis from pJ504 using o5161 et 5162 |
| peT43.1a | pBR322*::*T7 promoter::NusTag::His Tag | *ori*ColE1; [Carb$^R$]<br>*(41)* |
| pM413 | peT43.1a*::intIA$_{Vch}$* | *ori*ColE1; [Carb$^R$]<br>*(41)* |
| pM415 | peT43.1a*::intIA$_{Vch}$* (Y302F) | *ori*ColE1; [Carb$^R$]<br>*(41)* |
| pD060 | pSW23T*::attC$_{aadA7}$* (bs) | *oriV*$_{R6K\gamma}$, *oriT*$_{RP4}$; [Cm$^R$] *(10)* |
| p7848 (pMP7) | pSW23T*::araC* P$_{BAD}$-*ccdB* | *oriV*$_{R6K\gamma}$, *oriT*$_{RP4}$; [Cm$^R$] *(37)* |
| pK590 | pMP7*::*Δ*attIA* | *oriV*$_{R6K\gamma}$, *oriT*$_{RP4}$; [Cm$^R$] *(3)* |
| pP897 | pMP7*::TA4*STOP/BamH1* | *oriV*$_{R6K\gamma}$, *oriT*$_{RP4}$; [Cm$^R$]<br>Assembly PCR of two fragments of the toxin of TA4 (brnT) amplified with oligos o6482/o6485 and o6484/o6483 for replacement of a CAA into a TAA in the coding sequence. Cloning into a EcoRI/PstI digested pMP7. Then addition of a BamH1 site close to the newly added STOP codon by reverse PCR (o6498/o6499) for screening the successful allelic replacements. |
| pP898 | pMP7*::TA10*STOP/BamH1* | *oriV*$_{R6K\gamma}$, *oriT*$_{RP4}$; [Cm$^R$]<br>Idem for the toxin of TA10 (relE). Assembly with o6486/o6489 and o6488/o6487 amplified fragments. Reverse PCR with o6500/o6501. |
| pP900 | pMP7*::TA16*STOP/BamH1* | *oriV*$_{R6K\gamma}$, *oriT*$_{RP4}$; [Cm$^R$]<br>Idem for the toxin of TA16 (parE). Assembly with o6490/o6493 and o6492/o6491 amplified fragments. Reverse PCR with o6502/o6503. |
| pP902 | pMP7::*TA18*STOP/BamH1* | *oriV*$_{R6K\gamma}$, *oriT*$_{RP4}$; [Cm$^R$] |

| | | |
|---|---|---|
| | | Idem for the toxin of TA18 (VCA497). Assembly with o6494/o6497 and o6496/o6495 amplified fragments. Reverse PCR with o6504/o6505. |
| pQ880 | pELSA | *oripSC101*; [Rif$^R$]<br>Design of the ELSA according to Reis and coll *(28)*. Synthesis of the ELSA by Twist Bioscience. ELSA fragment amplified (o6581/o6582) and inserted by Gibson Assembly into the o6583/o6584 amplified pG086 plasmid. |
| pN700 | *dCas9-sgRNA* | *orip15A*; [Cm$^R$] *(38)*<br>This dCas9-sgRNA plasmid expression system (pFR48) was derived from plasmid pJF1 (a gift from Eligo Bioscience) harbouring a constitutively expressed sgRNA and *cas9* under the control of a DAPG-inducible PhlF promoter. |
| pP111 | *dCas9-PmCDA1/UGI/LVA-sgRNA* | *orip15A*; [Cm$^R$]<br>Fusion of dCas9 with Cytidine deaminase, Uracil glycosylase inhibitor and degradation tag as described by Banno and coll *(27)*.<br>*PmCDA1/UGI/LVA* fragment synthesized by Twist Bioscience, reamplified (o7023/o7026) and inserted by Gibson Assembly into o7024/o7025 amplified pN700. |
| pP388 | pP111::*lacZ* | *orip15A*; [Cm$^R$]<br>annealing of complementary o6435/o6436 primers, cloned into pP111 by Golden Gate Assembly |

**Table S3: Primers used in this study**

| Primers | Sequences |
|---|---|
| **Primers used for the Y302F integrase expression vector** | |
| 5161 | CCGATGTAAAAACGACACAGATTTTCACTCAGTTCTTGATAGAGG |
| 5162 | CCTCTATCAAGAACATGAGTGAAAATCTGTGTCGTTTTTACATCGG |
| **Primers used for *Vibrio cholerae* parental strain construction** | |
| 4286 | GCTGTAGCACGTATGCTTCC |
| 4302 | AAGGTAAGGGGGGTAAAAATCGCAC |
| 4657 | CTTCACCTATGACCCCTGAGCAAAT |
| 4662 | AGAAGTGCATGTTTCATCTCCCCCT |
| **Primers used for *LacZ* guide-containing construction** | |
| 6435 | AAACATCCGCAAATGGATATTGGA |
| 6436 | TAGTTCCAATATCCATTTGCGGAT |
| 3812 | CAAGTTGATAACGGACTAGCC |
| 2406 | CATCATGCCGTTTGTGATGG |
| **Primers used for Base editor construction** | |
| 6581 | TTGACATGAGCTCGTCGTCAGG |
| 6582 | TAATCAGGATACTTCTCACTGTAAGC |
| 6583 | TATCCTGACGACGAGCTCATGTCAAATAGTTCTCACTGAATATTTAACTGG |
| 6584 | CTTACAGTGAGAAGTATCCTGATTACATGGGTTCTTTGCGAAATCATGAG |
| 7023 | CTGAGTCAGTTGGGCGGTGAC |
| 7024 | GTCACCGCCCAACTGACTCAG |
| 7025 | CCCGCTTCGGCGGGTTTTTAC |
| 7026 | GTAAAAACCCGCCGAAGCGGG |
| 6370 | CCTCGCATTTTTCTCGTAATAGAG |
| 6371 | CGAAATCTTTAGCATTAGAAAAGTCGAAG |
| 6372 | CTGGAGTATTACGACCGTTACGAGATCAAACATCCACTAGTC |
| 6373 | GACTAGTGGATGTTTGATCTCGTAACGGTCGTAATACTCCAG |
| 6374 | CCTTACACTACGAGCGATAATTGACAGGTGAACGCTCAGC |
| 6375 | GCTGAGCGTTCACCTGTCAATTATCGCTCGTAGTGTAAGG |
| 6376 | GACTTCTGGTGCTACCCGATTGACATGAGCTCGTCGTCAG |
| 6377 | CTGACGACGAGCTCATGTCAATCGGGTAGCACCAGAAGTC |
| **Primers used for *dcas9* gene sequencing** | |
| 6342 | ATGGATAAGAAATACAGCATAGGC |
| 6343 | ATTTCCTGATCGAAGGCGACC |
| 6344 | CTTCTGAAAGCGCTGGTACG |
| 6345 | AAACCTGCCGAACGAAAAGG |
| 6346 | TTAACGGTATCCGTGACAAACAG |

| | |
|---|---|
| 6347 | TCCGATTACGACGTTGACCAC |
| 6348 | ATATCCGAAACTGGAAAGCGAATTTG |
| 6349 | CCATTATGGAACGTAGCTCTTTC |
| 6350 | AAACGTTACACCTCCACTAAAGAAG |
| 6351 | TTACGACGGGTGTAACGGCG |
| 6445 | AACGTCGTCGTGAGATGCCG |
| **Primers used for the construction of TA inactivated strains by allelic exchange** | |
| 6482 (TA4i) | GTATCGATAAGCTTGATATCGAATTCCTTTGATGCAACATTTATTCATTTC |
| 6483 (TA4i) | GTGGATCCCCCGGGCTGCAGCTCACACTTGAATGATGAGACAC |
| 6484 (TA4i) | CATACAGCTTAAGGATTGTGGAATG |
| 6485 (TA4i) | CATTCCACAATCCTTAAGCTGTATG |
| 6498 (TA4i) | GATCCTGATATTCGTACCTC |
| 6499 (TA4i) | CTCTCAGTTCGGCGTTCACG |
| 6486 (TA10i) | GTATCGATAAGCTTGATATCGAATTCGGCCGAATACAGAATATCTGTG |
| 6487 (TA10i) | GTGGATCCCCCGGGCTGCAGCAATTGCTTATAAATTTTATGCCTAACG |
| 6488 (TA10i) | CTTCTTGAACTACTGTTGGAGTGG |
| 6489 (TA10i) | CCACTCCAACAGTAGTTCAAGAAG |
| 6500 (TA10i) | GATCCACATGTTCCATCAGC |
| 6501 (TA10i) | CTCAAGGCGTTCTATTAGTTTC |
| 6490 (TA16i) | GTATCGATAAGCTTGATATCGAATTCGTATGTTCTTACCGATGAAAACGG |
| 6491 (TA16i) | GTGGATCCCCCGGGCTGCAGAGTTTGTTTGATGGACAGGGTTTTC |
| 6492 (TA16i) | CTAAACAACTGGTGTAAACCGTTTTTACAAAAG |
| 6493 (TA16i) | CTTTTGTAAAAACGGTTTACACCAGTTGTTTAG |
| 6502 (TA16i) | CCGCCCAGACTCTGGAAAATC |
| 6503 (TA16i) | ATCCCTCCAGAACTAGAACACC |
| 6494 (TA18i) | GTATCGATAAGCTTGATATCGAATTCTGGTCGGACTTTTAGTCTTGCGG |
| 6495 (TA18i) | GTGGATCCCCCGGGCTGCAGAGGAGCGATAAGCACTCGTACG |
| 6496 (TA18i) | CTTACCCAATAAGTAACGAAAACGTGCTG |
| 6497 (TA18i) | CAGCACGTTTTCGTTACTTATTGGGTAAG |
| 6504 (TA18i) | CTCACCATTCACCGCTTTATTC |
| 6505 (TA18i) | GATCCAGGTAAGTTAGGTGATTTATG |
| 6470 | CTGCAGCCCGGGGGATCCAC |
| 6471 | GAATTCGATATCAAGCTTATCGATAC |
| 775 | GCGGTGCTCAACGGGAATCC |
| 776 | AAAAATGCACCGGGGCCAGC |
| **Primers used for the TA inactivated sequencing** | |
| 6531 (TA1-7) | GGAAACAGCTATGACCATGATTACGAATTCATATGTAGTTCTCTTTGACTGAATTACACA |
| 6532 (TA1-7) | GCATGCCTGCAGGTCGACTCTAGAGGATCCTTAGTTTGCAATCGGAGACTAAGAAACAAAGCG |
| 6533 (TA2-14) | GGAAACAGCTATGACCATGATTACGAATTCAGATTTTGATGATTTTTGATTGAGAGAAAC |
| 6534 (TA2-14) | GCATGCCTGCAGGTCGACTCTAGAGGATCCTTAGTTTGCTAAAGGTAGTGGATAACTAGCC |
| 6535 (TA3-19) | GGAAACAGCTATGACCATGATTACGAATTCCACAACCTACTGATTTTTATGGATTAAGCA |
| 6536 (TA3-19) | GCATGCCTGCAGGTCGACTCTAGAGGATCCTTATGTGCACAATTGACTCATTAGAATAAATC |

| | |
|---|---|
| 6537 (TA4) | GGAAACAGCTATGACCATGATTACGAATTCATACGAAAAGTTTGGAGGATAAATATCACC |
| 6538 (TA4) | GCATGCCTGCAGGTCGACTCTAGAGGATCCTTAGCCGTTTATCTCACACTTGAATGATGAGACACG |
| 6539 (TA5) | GGAAACAGCTATGACCATGATTACGAATTCAAGATAGCTCTCATGATATCTCCAAATACC |
| 6540 (TA5) | GCATGCCTGCAGGTCGACTCTAGAGGATCCTTAGTTGCCAATCAGCAACAAATCAGTCGTTTGG |
| 6541 (TA6) | GGAAACAGCTATGACCATGATTACGAATTCCTAATATTCCTAAATGTTTGAAATGGCACA |
| 6542 (TA6) | GCATGCCTGCAGGTCGACTCTAGAGGATCCTTATGCGCCGAATATTGGGTTCACATCCATGC |
| 6543 (TA8) | GGAAACAGCTATGACCATGATTACGAATTCAAATCATTGTTAAACAATGGGCTACGAAGC |
| 6544 (TA8) | GCATGCCTGCAGGTCGACTCTAGAGGATCCTTATGTACGTGCGCTTTGTTCCAGCGAAACAAAC |
| 6545 (TA9) | GGAAACAGCTATGACCATGATTACGAATTCGAAAGAGTGCGTTATGTAACCAGTGTGTGC |
| 6546 (TA9) | GCATGCCTGCAGGTCGACTCTAGAGGATCCCAATCACAACCTGCTGATTTTTTATGGATTAAGCA |
| 6547 (TA10) | GGAAACAGCTATGACCATGATTACGAATTCTATCAAGCAATTGCTTATAAATTTTATGCC |
| 6548 (TA10) | GCATGCCTGCAGGTCGACTCTAGAGGATCCTTAAACTTCATCAGTCGTCTAGCCGTTGCAGC |
| 6549 (TA11) | GGAAACAGCTATGACCATGATTACGAATTCATGGAAGGGTGGGCATCAGAAACACACACA |
| 6550 (TA11) | GCATGCCTGCAGGTCGACTCTAGAGGATCCTTATAGCTCGGCTATGTGTGACAACGTTTCTGG |
| 6551 (TA12) | GGAAACAGCTATGACCATGATTACGAATTCTTAGATTTGCTTTTAGCGGAAAGTGGCTCA |
| 6552 (TA12) | GCATGCCTGCAGGTCGACTCTAGAGGATCCTTATGTTTAGAGAGCTGAATTTTGTTCTTTTGCG |
| 6553 (TA13) | GGAAACAGCTATGACCATGATTACGAATTCACTGAACAAGGCTCACAAATAGCACGCATA |
| 6554 (TA13) | GCATGCCTGCAGGTCGACTCTAGAGGATCCTTGTGCCGCTAACCGAAGCGTGAAGCCAGTT |
| 7027 (TA15) | GGAAACAGCTATGACCATGATTACGAATTCCTAGAACAAGAGCAATCATCAGCACACACA |
| 7028 (TA15) | GCATGCCTGCAGGTCGACTCTAGAGGATCCTTATATTCGTAGCCTATTGATTAAAAAGTTGTTC |
| 6557 (TA16) | GGAAACAGCTATGACCATGATTACGAATTCGTAGTGGCGTTGGCTGCCCCTTAAGCGGGC |
| 6558 (TA16) | GCATGCCTGCAGGTCGACTCTAGAGGATCCTTAGCCGTTTTTTTCAAACCTACTGTTTCG |
| 6559 (TA17) | GGAAACAGCTATGACCATGATTACGAATTCTATTACTACATGAGCTTCGTAGCACAAACC |
| 6563 (TA17) | GCATGCCTGCAGGTCGACTCTAGAGGATCCTTAGTTTTGGGCGCTAGGAAGAAAATTTATCTA |
| 6561 (TA18) | GGAAACAGCTATGACCATGATTACGAATTCGAAAGAGTAGCCGTTTCAGAACCACAAACA |
| 6562 (TA18) | GCATGCCTGCAGGTCGACTCTAGAGGATCCTTATATGCGTGGTTCCTTGTTTGCCACTGTTC |
| **Primers used to confirm the recombination events** | |
| 5778 | GTCAAGAGGCTATACAGACATCAGC |
| SWend | CCTCACTAAAGGGAACAAAAGCTG |
| 1401 | GATGGCAACATATTTAACTCATCTTGAGG |
| **Primers used for the electromobility shift assay** | |
| VCR | CCGGTTATAACGCCCGCCTAAGGGGCTGACAACGCGAAGCGTTGACAGTCCCTCTT GAGGCGTTTGTTATAACCGG |

#### Supplementary movies

**Movie S1:** Time-lapse movie of SCI Reinverted strain growth

a. Time-lapse movie of cells (SCI Reinverted strain) growing onto agarose pad in the presence of arabinose to induce the expression of IntIA. The growth of single cells into microcolonies was followed for 170 min. Time and scale bar are indicated.
b. As in a. but focusing on three microcolonies.

**Movie S2:** Time-lapse movie of SCI Inverted strain growth

a. Time-lapse movie of cells (SCI Inverted strain) growing onto agarose pad in the presence of arabinose to induce the expression of IntIA. The growth of single cells into microcolonies was followed for 170 min. Time and scale bar are indicated.
b. As in a. but focusing on three microcolonies.
c. As in a. but focusing on cell lysis.

Events of cell lysis are displayed, characterized by a sudden release of diffracting material.
